# A novel phyllosphere resident *Protomyces* species that interacts with the *Arabidopsis* immune system

**DOI:** 10.1101/594028

**Authors:** Kai Wang, Timo Sipilä, Sitaram Rajaraman, Omid Safronov, Pia Laine, Agate Auzane, Alfredo Mari, Petri Auvinen, Lars Paulin, Eric Kemen, Jarkko Salojärvi, Kirk Overmyer

**Affiliations:** Organismal and Evolutionary Biology Research Program, Faculty of Biological and Environmental Sciences, and Viikki Plant Science Centre; Institute of Biotechnology, University of Helsinki, P.O. Box 65 (Viikinkaari 1), FI-00014 Helsinki, Finland; Interfaculty Institute of Microbiology and Infection Medicine Tübingen, IMITP, University of Tübingen, 72076 Tübingen, Germany; School of Biological Sciences, Nanyang Technological University, 60 Nanyang Drive, Singapore 637551, Singapore

## Abstract

We describe the genome contents of six *Protomyces* spp. that are pathogenic within the typical host range of the genus and a novel *Protomyces* strain (SC29) that was previously isolated from the phylloplane of wild *Arabidopsis thaliana* (*Arabidopsis*), an atypical or possible alternate host. Genome-wide phylogenetic analysis defined SC29 as a distinct *Protomyces* sp. Analysis of gene family expansions, gene retention, and gene loss patterns among these *Protomyces* spp. lead us to hypothesize that SC29 may have undergone a host jump. The role of phyllosphere residency in the lifecycle of *Protomyces* spp. was previously unknown. Genomic changes in SC29 and all other *Protomyces* spp. were consistent with adaptations to the plant phylloplane. As predicted by our analysis of its mating locus, SC29 did not cause disease on *Arabidopsis* as a single strain, but could persist in its phylloplane, while the closely related *P. inouyei* does not. SC29 treated *Arabidopsis* exhibited enhanced immunity against *Botrytis cinerea* infection, associated with activation of MAPK3/6, camalexin, and SA-signalling pathways. We conclude that SC29 is a novel *Protomyces* sp. able to survive in the *Arabidopsis* phylloplane and that phylloplane residency is an important element in the lifecycle of *Protomyces* spp.

## INTRODUCTION

*Protomyces* spp. are yeast-like members of the subphylum Taphrinomycotina, which retain many ancestral characteristics and are known phytopathogens, causing disease on host plants within the Compositae (Asteraceae) and Umbelliferae (Apiaceae) families (1, 2). Species in *Protomyces* and its sister genus *Taphrina* (order Taphrinales) share similar dimorphic lifestyles, possessing both a haploid budding yeast phase and invading their hosts in their hyphal form (3). Hyphae of species in the *Taphrina* are dikaryotic but diploid in *Protomyces* (1, 4), illustrating also marked differences in lifecycles exist between members of these two genera. A common virulence strategy is shared, both cause plant tumours or gall-symptoms and are well known to produce plant hormones (2, 3). As members of the Taphrinomycotina with notable ancestral characteristics and similarities to basidiomycete yeast-like fungi, species in the genera *Protomyces* and *Taphrina* are of considerable phylogenetic and evolutionary interest. However, these fungi cause diseases primarily on wild and woody species and remain understudied (4). Recent reference genome assemblies have opened these taxa to molecular evolutionary studies, increasing interest and proving the value of non-model species in the Taphrinomycotina (5-8).

The *Protomyces* also have scientific value as plant associated yeasts. Yeasts have a spectrum of relationships with their host plants ranging from the pathogens to growth-promoting symbionts. Our understanding of plant associated yeasts lags behind that of other classes of microbes. This is the case for *Arabidopsis thaliana* (*Arabidopsis*), a model plant widely used when studying the molecular basis of plant-microbe interactions (9). Currently, no well-characterized systems for the study of *Arabidopsis*-yeast interactions exist. Several yeasts have been identified as hub taxa, which restructure microbiome community composition when present in the *Arabidopsis* phyllosphere (10, 11). The phyllosphere is gaining attention as a microbial habitat, with focus primarily on prokaryotic residents. Eukaryotic resident species are now being isolated, but their genomic adaptations to the phyllosphere remain underexplored. Phyllosphere residency in the yeast form is an important component of the lifecycle of *Taphrina* species; however, its significance in the lifecycle of *Protomyces* species remains unknown (2, 3).

We have previously isolated a novel strain of *Protomyces* that is associated with *Arabidopsis* (12, 13). Here we characterize its interactions with *Arabidopsis* and compare the genome to six reference species of *Protomyces.* We present evidence of a new species that is able to survive in the phylloplane environment of *Arabidopsis.* Further, all *Protomyces* spp. examined here bear genomic adaptations to the phyllosphere supporting the importance of this living space in the lifecycle of *Protomyces* species.

## MATERIALS AND METHODS

### Protomyces strains and culture

*Protomyces* reference species (Table 1) were obtained from the ARS culture collection (https://nrrl.ncaur.usda.gov/). SC29 was isolated from wild *Arabidopsis* (13). All species were purified twice from single colonies and cultured on glucose, yeast-extract, peptone (GYP) agar at 21°C, unless otherwise indicated.

**Table 1.** Genome sequencing, assemblies and annotation statistics of seven *Protomyces* species in this study. * Gene set completeness was estimated using BUSCO. ^‡^ Reference strains were obtained from the ARS culture collection (https://nrrl.ncaur.usda.gov/).

| Species | *P. sp. SC29* | *P. gravidus* | *P. inouyei* | *P. inundatus* | *P. lactucaedebilis* | *P. macrosporus* | *P. pachydermus* |
| --- | --- | --- | --- | --- | --- | --- | --- |
| Abbreviation | SC29 | Pgra | Pino | Pinu | Plac | Pmac | Ppac |
| Strain<sup>†</sup> | C29 | Y-17093 | YB-4354 | Y-6349 | YB-4353 | Y-1287 | YB-4355 |
| Host family | ? | Compositae | Compositae | Umbelliferae | Compositae | Umbelliferae | Compositae |
| No. known hosts | ? | 2 | 1 | 3 | 1 | 27 | 7 |
| Bioproject | PRJNA486901 | PRJNA486711 | PRJNA486715 | PRJNA486783 | PRJNA486789 | PRJNA486796 | PRJNA486895 |
| Biosample | SAMN09867691 | SAMN09862461 | SAMN09862466 | SAMN09863624 | SAMN09863782 | SAMN09864018 | SAMN09867611 |
| SRA | SRR8109439 | SRR7725366 | SRR7725445 | SRR7725484 | SRR7725555 | SRR7725730 | SRR7725731 |
| Genome | QXMI00000000 | QXDP00000000 | QXDQ00000000 | QXDR00000000 | QXDS00000000 | QXDT00000000 | QXDU00000000 |
| No. of contigs | 149 | 222 | 353 | 370 | 329 | 146 | 252 |
| Largest contig | 877.8 kb | 685.7 kb | 604.9 kb | 460.7 kb | 443.5 kb | 986.12 kb | 498.7 kb |
| Assembly size | 11.9 Mb | 11.5 Mb | 12.9 Mb | 14. 1 Mb | 12.8 Mb | 13.1 Mb | 12.0 Mb |
| GC (%) | 50.9 | 50.9 | 51.1 | 52.8 | 51.1 | 52.7 | 51.2 |
| Scaffold *N50* | 449.7 kb | 228.1 kb | 213.7 kb | 132.7 kb | 181.4 kb | 327.6 kb | 294.8 kb |
| Scaffold *N75* | 233.7 kb | 127.0 kb | 122.0 kb | 77.1 kb | 94.0 kb | 202.4 kb | 161.1 kb |
| Scaffold *L50* | 10 | 15 | 20 | 29 | 24 | 14 | 16 |
| Scaffold *L75* | 20 | 32 | 39 | 65 | 47 | 26 | 31 |
| N's per 100 kp | 0 | 0.02 | 0 | 0,01 | 0 | 0.01 | 0.07 |
| Annotated genes | 5514 | 5422 | 5956 | 6287 | 5952 | 5912 | 5612 |
| Completeness * | 86.0% | 85.1% | 86.1% | 88.2% | 86.3% | 87.0% | 86.2% |
| ANI to SC29 | 100 | 78 | 86 | 78 | 86 | 78 | 85 |
| AAI to SC29 | 100 | 74 | 91 | 72 | 91 | 72 | 90 |

### SC29 pre-treatment, Botrytis infection, and qPCR

Live and autoclaved SC29 were harvested, washed, and suspended at 1 OD_600_ (7 × 10^6^ cfu/ml) in sterile water. Three-week-old soil-grown *Arabidopsis* were sprayed with one of the suspensions (0.2 ml/plant), or water as a control, then infected with a *Botrytis* spore suspension (14) three days after SC29 treatments. Plants were grown in a chamber with 12/12h (light/dark), 23/18°C, and 65/75% relative humidity. Pre-treated and infected plants were collected at 30 and 60 min for western blot, and at 24 and 72 h for qPCR. Four of five plants were pooled, frozen in liquid nitrogen, and stored in −80°C. Leaf lesions were photographed at 72 h and diameters measured in ImageJ. Lesion size data were statistically analysed with scripts in R (version 3.5.1) using the nlme package, a linear mixed model with fixed effect for treatment was fitted to the data with a random effect for biological repeat. The model contrasts were estimated with multcomp package, and the estimated p-values were subjected to single-step correction. RNA isolation and qPCR were performed as previously described (14). Primer sequences, amplification efficiencies, and reference genes are listed in file S2. Raw cycle threshold values were analysed with Qbase+ 2.1 (15). Significance was estimated by glht package in R (3.5.1) using false discovery rate adjustment of p-values.

### Protomyces growth assays

Three-day-old cultures of *P. inouyei* (OD_600_ = 0.14, 5 µl/leaf) and SC29 (OD_600_ = 0.10, 5 µl/leaf) were drop inoculated on the adaxial side of 24-day-old *Arabidopsis* leaves, aseptically grown on 0.5xMS agar plates. Sterile foil on an agar plate was used as a control for non-specific surface growth. Growth in the phylloplane was assayed by re-isolating *Protomyces* from inoculated leaves by placing leaves in 2 ml Eppendorf tubes with 1 ml 0.025% Silwet-L77 in water and shaking (800 rpm; VWR microplate shaker) for 1 hour, after which 5 µl wash solution was serial diluted and pated on GYP agar for colony counting. Pooled data from five biological repeats were analysed using a linear mixed model in R, as above. Plants for long term field infections were sown and germinated in September in a covered cage under ambient conditions using seeds of the accessions Col-0 and a line Kivikko (Kvk1) locally collected at the site where yeast isolation samples were collected. Seven-day-old seedlings were transplanted 5 per pot in 10 cm square plastic pots with a 1:1 mix of peat and vermiculite and further grown under the covered cage. Pots containing two-week-old plants were buried in sand in the Helsinki University Viikki campus experimental field in a triplicate block design at three different sites in the field and spray inoculated with water (mock) or SC29 (OD_600_ = 0.1 and 1.0, ca. 1 ml per pot). Plants remained in the field over the course of the next seven months (until early May), with snow cover for about four months, and were visually examined and photographed at various times. In May, final photos and visual examinations were done and representative Mock control and infected plants were sampled for yeast isolations, performed as previously described (13).

### Presence of Protomyces OTUs in environmental sequencing experiments

Occurrence of *Protomyces* OTUs on the *Arabidopsis* Columbia (Col-0), Keswick (Ksk1), San Feliu (Sf-1), and Wassilewskija (Ws-0) accessions was monitored by mining ITS barcode library sequencing data from common garden field planting experiments repeated in the years 2014-2016. Experiments were performed in Cologne, Germany (N Coord. 50° 57’ 21.56” E Coord. 6° 51’ 40.20”). Accounting for host genetic variation, four *Arabidopsis* genotypes were chosen based on their variation in susceptibility towards *A. laibachii* and ability to survive within the field conditions (10). Surface sterilized seeds were stored at 4°C in 0.1% agarose for one-week prior sowing on water soaked jiffy pots (www.jiffypot.com) and kept for two weeks in a glasshouse (10 h light, 14 h darkness, 23/20 °C, 60% humidity). Plants were then colour coded and planted across the plot at random. Samples for sequencing ITS1 libraries were collected in triplicate monthly, one per subplot, over the course of the *Arabidopsis* growing season (November to March). Sample handling, library preparation, and sequencing with the Illumina MIseq platform are as previously described in Agler et al. (10).

The presence of *Protomyces* species in publicly available environmental metagenomic barcode marker sequencing data was determined by homology searches with BLASTn in August 2019 with default search parameters and using the NR database including environmental data. ITS sequences from SC29 and the six reference *Protomyces* species were used as queries. Results from BLAST searches were manually tallied and summarized in Figure S2b.

### Genomic DNA extraction

Genomic DNA was isolated from cultures of the haploid yeast form for each species using a chromosomal DNA extraction method (16). Bead beating was done using a vortex (Vortex genie 2, Scientific Industries, Inc., United States) with multiple 2 ml tube adapter (2 min, full speed). Beads (0.2-0.3 g/tube) with 0.75-1.00 mm dimeter were used (Retsch GmbH, Germany). The quantity of isolated DNA was estimated using Nanodrop ND-1000 (Thermo Scientific, USA) and qubit fluorometer (Thermo Fisher Scientific, USA). The DNA yield was in average 23.9 µg/strain with standard deviation 12.6 µg assayed with qubit fluorometer. ITS fragments were amplified and sequenced as previously described (13) to confirm the species identity prior to sequencing.

### Genome assemblies and annotations

Genome sequencing was done at the DNA Sequencing and Genomics Laboratory, Institute of Biotechnology, University of Helsinki. MiSeq Sequencing System (Illumina, California USA) was used to sequence pair end reads with average read length 231 bp with standard deviation 4.8 bp. Paired end sequencing libraries were prepared using Paired-End Sample Preparation Guide (Illumina). The four partial Illumina sequencing runs produced 758,373 read pairs in average (Standard deviation 17,341 read pairs). These reads were assembled using SPAdes v. 3.1.1 (17).

The *Protomyces* genome assemblies were annotated using the Hidden Markov Model based gene predictor Augustus version 2.5.5 (18). The gene predictor was trained using RNASeq data obtained from *Taphrina betulina* (Bioproject: PRJNA188318) (19). Paired RNA raw reads (Run 1) were aligned against the *Taphrina betulina* genome to generate the BAM alignment file using TopHat version 2.0.11 which internally used Bowtie version 2.1.0.0 (20). The unpaired RNA raw reads (Run 2) (21) were also aligned against the genome using TopHat using junction information from the paired read alignment. The BAM alignment files coming from Run 1 and Run 2 were merged using Samtools version 0.1.19-44428cd (22). Transcripts were then generated from the merged BAM alignment file using Cufflinks version 2.2.1 (23). These transcripts were then splice-aligned against the *Taphrina betulina* genome using PASA version r20130605p1 (24) to generate complete ORFs. These ORFs combined with 950 manually annotated genes and CEGMA (Core Eukaryotic Genes Mapping Approach) genes (25) were used to train Augustus. The gene set completeness per genome was estimated using BUSCO (Benchmarking Universal Single-Copy Orthologs) version 3.0.2 (26) internally using the Ascomycota odb9 fungal dataset. The predicted gene models were assigned human-readable functional descriptions using the tool AHRD (Automated Assignment of Human Readable Descriptions) version 3.3.3 (https://github.com/groupschoof/AHRD). Conserved protein domains were defined using HMMER (v3.2.1; www.hmmer.org; with the parameters tblout, E value 1e-30) with the Pfam database (File S1) (27). The gene models were initially aligned against two custom BLAST databases containing fungal proteins from SWISSPROT and TREMBL respectively. The BLAST results along with GO annotation information of the fungal proteins in UniProt format were then passed to the AHRD tool which parses the results and assigns the best description and GO information to the gene models. Gene family evolutionary rate and ancestral size estimation of the orthogroups were carried out using badirate v.1.35 (28). A species-level rooted phylogenetic tree and orthogroup size information were provided as input for the badirate software. The BDI turnover rate model (Birth, Death and Innovation) was selected. The FR branch model (Free Rates) was used which assumes that every branch in the phylogenetic tree has its own turnover rate. Maximum Likelihood statistical method was used to estimate the turnover rates. The outlier branches reported per orthogroup, which did not evolve under the estimated turnover rates, if any, were identified as significant branches for further analysis.

For genome synteny analysis, whole genome alignment was performed with Mummer 4.0.0beta2 (nucmer) with default settings. Mummerplot (parameter --fat) was applied to shape the optimal co-linear order of contigs. Mummerplot output ps files were edited with CorelDRAW2018.

### Identification and analysis of mating loci

Mating (MAT) locus genes were identified by homology searching utilizing local BLASTp (29)) against annotated proteins from *Protomyces* species. Searches with tBLASTn were also used to query *Protomyces* genome sequences for additional genes that were not annotated. *T. deformans* and *S. pombe* MAT loci proteins (encoded by *matMc, matPi*, and *matPc*), and proteins encoded by MAT loci associated genes (*sui1, rga7, cox10*), were used as search queries. Our analysis focused on those proteins that were similar to known mat proteins, correctly organized in loci with other mat proteins and found in close proximity to other *mat* loci associated proteins (i.e. *sui1, rga7, cox10*). Additionally, all HMG-box and homeodomain proteins were identified in *Protomyces* species and phylogenetic trees constructed with a collection of respective genes from *S. pombe* to ensure that the correct orthologs had been identified (not shown). Once *mat* loci were identified, the above phylogenetic trees, additional BLAST searches using the species own mat proteins against its own genome and annotated proteins, and protein alignments were used to rule out the possibility of additional silent MAT loci, consistent with a MAT switching (secondary homothalism). The *matMi* peptides, which are poorly conserved between even closely related species, were identified from six frame translations (EMBOSS Sixpack; www.ebi.ac.uk/Tools/st/emboss_sixpack/) of contigs bearing *matMc* based their size, proximity to *matMc*, and opposite orientation to *matMc*, as in (30). Contigs with *mat* loci were visualized using R package genoPlotR. Phylogenetic trees of mat proteins from *Protomyces* spp., including previously identified mat proteins from *S. pombe, Pneumocystis carinii, Pn jirovecii*, and *T. deformans* were constructed using methods described in (12).

### Genome-wide phylogenetic analysis

Orthologous protein families from 15 yeast species were inferred with OrthoFinder (v2.2.6; with default settings) (31) utilizing the Guidance2 algorithm (32) multiple sequence alignment. Totally, 1035 single-copy proteins with confidence scores >= 0.8 were selected and concatenated with the FASconCAT_v1.0 script (33). A Maximum likelihood phylogeny tree was built by RAxML v8 (34) with the rapid bootstrap algorithm (100 bootstraps). The tree was viewed and edited in iTOL (35). ANI and AAI values were estimated by genome-based distance matrix calculator (36). Phylogenetic tree with ITS marker were constructed using methods described in (12).

### Selection and analysis of CSSPs

Automated annotation did not capture candidate effector-like proteins (CELPs); thus a manual screen was used. We identified CELPs by defining small secreted proteins encoded in fungal genomes as follows. Open reading frames (ORFs) between 80 to 333 amino acids long were chosen and filtered for a secretion signal with SignalP 4.1 (37) to defines short secreted proteins (SSPs). After removing signal peptides, cysteines were counted; SSPs with ≥ 4 cysteines were regarded as cysteine-rich SSPs (CSSPs). OrthoVenn (38), orthofinder v2.2.6 (31), and Badirate (28) were used to analyse gene family evolution in CELPs. Known virulence factors were searched from SSPs and CSSPs using PHI base (39). Pfam domains of genomic proteins and SSPs were searched with HMMER v3.2.1 (http://hmmer.org/) using hmmscan (tblout, E value 1e-30).

### Genomic mining for multiple enzymes of important traits

Genome assemblies and annotations of all seven *Protomyces* species were applied as local database. Fungal HHK (hybrid histidine kinases) protein sequences were collected from prominent plant-pathogenic fungi (40) for BLASTp (E value < 1e^-30^) (29). Proteins involved in carotenoid biosynthesis (File S10) were collected from NCBI and UniProt and used as queries in BLASTp (E value <1e^-30^) (29). Sequences of enzymes involved in IAA synthesis of *T. deformans* (File S11) were selected as reference queries for tBLASTn (E value <1e^-5^, score >100 and identity >50%) (29). Duplicate hits were manually removed for all BLAST searches. Carbohydrate-active enzymes (CAZymes) in *Protomyces* genomes were searched with dbCAN (41).

### Indolic-compound production, and auxin-activity assays

Sterile filtrates (0.20 µm filter) from cultures on, GYP, GYP + 0.1% tryptophan, or nitrogen base with glucose, were tested for indolic compound production, as previously described (13, 42). Statistics were performed with *t*-test in R 3.5.1. Sterile seeds of *Arabidopsis thaliana* (arabidopsis) bearing the artificial auxin-responsive promoter∷reporter (DR5∷GUS) fusion, were grown in 0.5xMS agar (1% sucrose, pH 5.7) for seven days under long-day conditions at 23°C. Staining followed previous protocol (13). Root hair phenotypes were captured using a LEICA MZ10F microscope with camera LEICA DFC490 24h after treatment of five-day-old arabidopsis seedlings with culture supernatant, as above.

### Protein extraction, SDS-PAGE, and Western blotting

Total protein was isolated from frozen rosettes using 100 mg plant material in 100 µl Lacus buffer (43). Protein concentration determined by Bradford assay (www.bio-rad.com/) with BSA as standard. Total protein (100 µg) was loaded in SDS-PAGE, transferred to a PVDF (immobilon–FL) membrane, and scanned with LI-COR Odyssey scanner. Equal loading was examined by amido black staining. Primary anti-TEpY (Phospho-p44/42 MAPK (Erk1/2) Thr202/Tyr204; CST®; www.cellsignal.com) rabbit monoclonal antibody specific for the activated form of MAPKs was used at a 1:2000 dilution and the secondary anti-rabbit IgG antibody (IRDye; www.abcam.com; 800CW Goat anti-Rabbit) at 1:10,000.

### Sequence accession numbers

All sequence data has been submitted under the following GenBank accession numbers: For ITS sequences: *Protomyces* sp. strain SC29 (SC29), LT602858, *P. gravidus* (Pgra), MK937055; *P. inouyei* (Pino), MK937056; *P. inundatus* (Pinu), MK937057; *P. lactucaedebilis* (Plac), MK937058; *P. macrosporus* (Pmac), MK937059; *P. pachydermus* (Ppac), MK937060 The GenBank accession numbers of assembled and raw genome sequencing data for all *Protomyces* spp. are found in table 1. Genome annotations available at genomevolution.org/coge/GenomeInfo.pl? with the following genome IDs: SC29, 53653; Pgra, 53651; Pino, 53654; Pinu, 53676; Plac, 54947; Pmac, 53670; Ppac 54948. The strain SC29 has been deposited in the University of Helsinki Microbial Domain Biological Resource Centre (HAMBI) Culture Collection (accession no. HAMBI3697) and the Collection of Microorganisms and Cell Cultures (DSMZ) culture collection (accession no. DSM 110145).

## RESULTS

### Novel Protomyces species of Arabidopsis

We previously isolated and characterized phylloplane yeasts from wild *Arabidopsis* growing in Helsinki, Finland (13). This included strains in OTU1, which was subsequently identified as a putative new *Protomyces* species based on ITS sequence (Genbank acc. no. LT602858) similarity (96%) to *P. inouyei* (13). Strain C29 (SC29) was selected for characterization in comparison to six species, which represent all species within the genus with strains available from yeast culture collections (Table 1). Reference species used and their abbreviations were; *P. gravidus* (Pgra), *P. inouyei* (Pino), *P. inundatus* (Pinu), *P. lactucaedebilis* (Plac), *P. macrosporus* (Pmac) and *P. pachydermus* (Ppac). To resolve the relationships within the genus *Protomyces*, we sequenced the genomes of these species. For genome statistics, conserved protein domains, and conserved synteny see (Table 1, File S1, Fig. S1). Phylogenetic analysis was conducted with whole genome data using 1035 single-copy proteins (Fig. 1). This analysis places SC29 as a distinct species within the *Protomyces*, as an outgroup to the clade composed of Ppac, Plac, and Pino, and as sister clade to Pgra, which are all pathogenic on Compositae family hosts. Pmac and Pinu form another distinct clade (Fig. 1) and infect Umbelliferae family hosts (Table 1). The same result was obtained using both the maximum likelihood (Fig. 1) and Bayesian inference (Fig. S2) methods. Further analysis describing and naming SC29 as a novel *Protomyces* species has been presented elsewhere (12).

**Figure 1.**
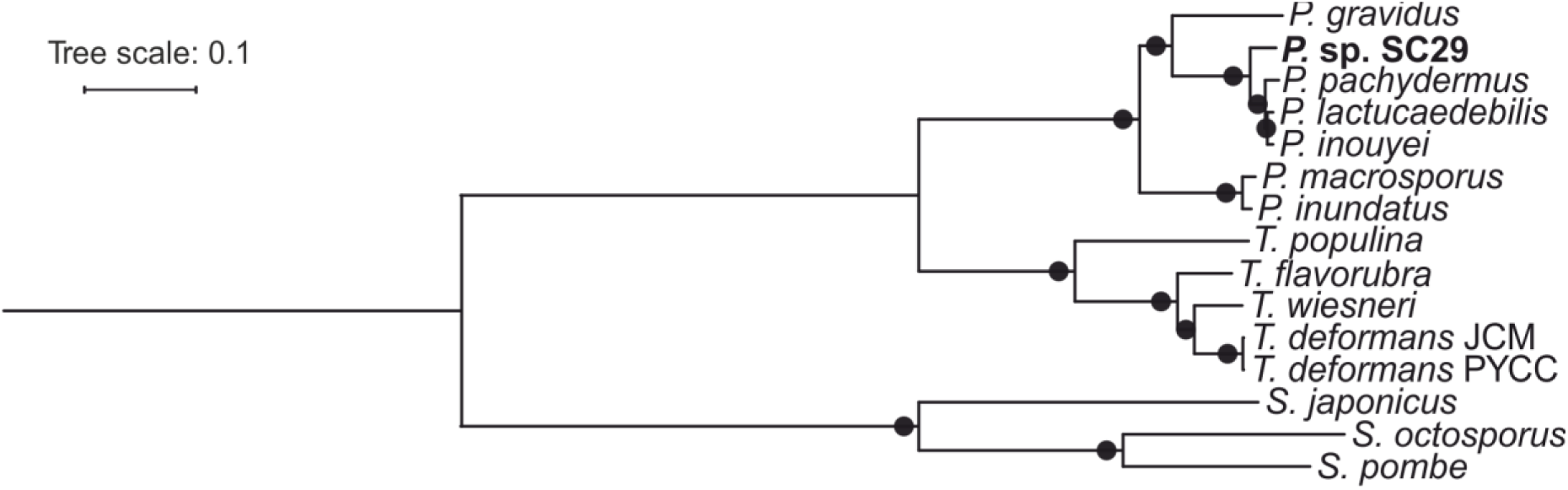
Phylogenetics of *Protomyces* species sequenced in this study. Species and abbreviations used are: *Protomyces* strain C29 isolated from wild *Arabidopsis* (SC29), *P. gravidus* (Pgra), *P. inouyei* (Pino), *P. inundatus* (Pinu), *P. lactucaedebilis* (Plac), *P. macrosporus* (Pmac) and *P. pachydermus* (Ppac). a) Whole genome phylogenetic tree of *Protomyces* and *Taphrina* spp. with three *Schizosaccharomyces* spp. used as an outgroup. RAxML and rapid bootstrapping (100x) were chosen for constructing the tree utilizing 1035 concatenated single-copy conserved protein sequences. Alignment quality control was achieved by applying sequence scores >= 0.8 in MAFFT analysis using Guidance2. Multiple aligned sequences of each species/strain were concatenated using FASconCAT_V1.0 and bootstrap values are indicated at the nodes, black circles denote 100% support. For whole genome alignment (synteny) data see Figure S3.

Our four strains of *Protomyces* were isolated from two independent *Arabidopsis* samples in Helsinki, Finland (13). To gain further evidence that SC29-like *Protomyces* OTUs are associated with *Arabidopsis*, we queried Illumina MIseq amplicon sequencing data targeting ITS1 from a three-year *Arabidopsis* garden time-course experiment in Cologne, Germany. This revealed the persistent presence of *Arabidopsis*-associated *Protomyces* OTUs on multiple *Arabidopsis* accessions, in varied abundance across the growth season (Fig. S3a). Phylogenetic analysis of ITS1 sequences representative of the OTUs found in this experiment indicated the many of them where related to SC29 (Fig. S3b). These data suggest SC29 is a distinct and novel species in the genus *Protomyces* and yeasts of the genus *Protomyces* are reproducibly associated with *Arabidopsis*.

To test if *Protomyces* species are found on plants other than their hosts, or in other environments, we queried publicly available environmental sequencing data (Fig. S2b, c). This revealed that *Protomyces* spp. are rarely found in environmental sequencing experiments, with few or no hits (Fig. S2c); however, SC29 as well as the closely related Pino and Plac had six, two, and two hits, respectively. An ITS phylogeny of the reference species and the six SC29 hits revealed uncultured strains highly similar to SC29 that have been found on broad bean (*Vicia faba*; clone 67_NA3_P32_F4; genbank KC965451.1) and in arctic soil (RP239-5; genbank KX067824.1) (44). This suggests that *Protomyces* spp. are largely restricted to their hosts and that SC29 and related strains may be more flexible in their environmental requirements.

### The mating loci of Protomyces species

All examined species of *Protomyces* are heterothallic (1, 2), i.e. they require conjugation with a strain of the opposite mating type to enter the plant pathogenic dikaryotic hyphal state, which initiates the sexual cycle. To address control of lifecycle transition, we identified and analysed the mating (MAT) loci (Fig. S4a-d; Files S2 and S3) in the sequenced genomes of *Protomyces* spp. revealing a configuration most consistent with the expected heterothallism, in most cases (Figure S4e-f). Each species had a single MAT locus either *matM* (Pmac, Pino, and Plac) or *matP* (Pgra, SC29, Ppac), with the exception of Pinu, which had both a *matM* and *matP*, suggesting it has a primary homothallic sexual cycle.

### SC29 persists on Arabidopsis phylloplane

Heterothallism in SC29 predicts that it should not be able to infect Arabidopsis as a single strain. To confirm this we tested many *Arabidopsis*-infection protocols, including chamber experiments at low temperatures and long-term field infections, mimicking the natural growth conditions of wild *Arabidopsis* (Table S1). No visible disease symptoms were observed. However, in field infections, SC29 survived overwinter on *Arabidopsis* and was re-isolated from infected plants the following spring. The growth of SC29 and Pino was assayed on the leaf surface of soil grown (not shown) and *in vitro* aseptic *Arabidopsis* cultures (Fig. 2a). In these experiments, SC29 persisted on the *Arabidopsis* leaf surface. In contrast, its closest relative, Pino, was unable to survive, demonstrating that adaptation to the *Arabidopsis* phyllosphere was specific to SC29. Further, SC29 was unable to survive on sterile foil placed on top of agar media plates, indicating it was not generally adapted to survive on surfaces without specificity.

**Figure 2.**
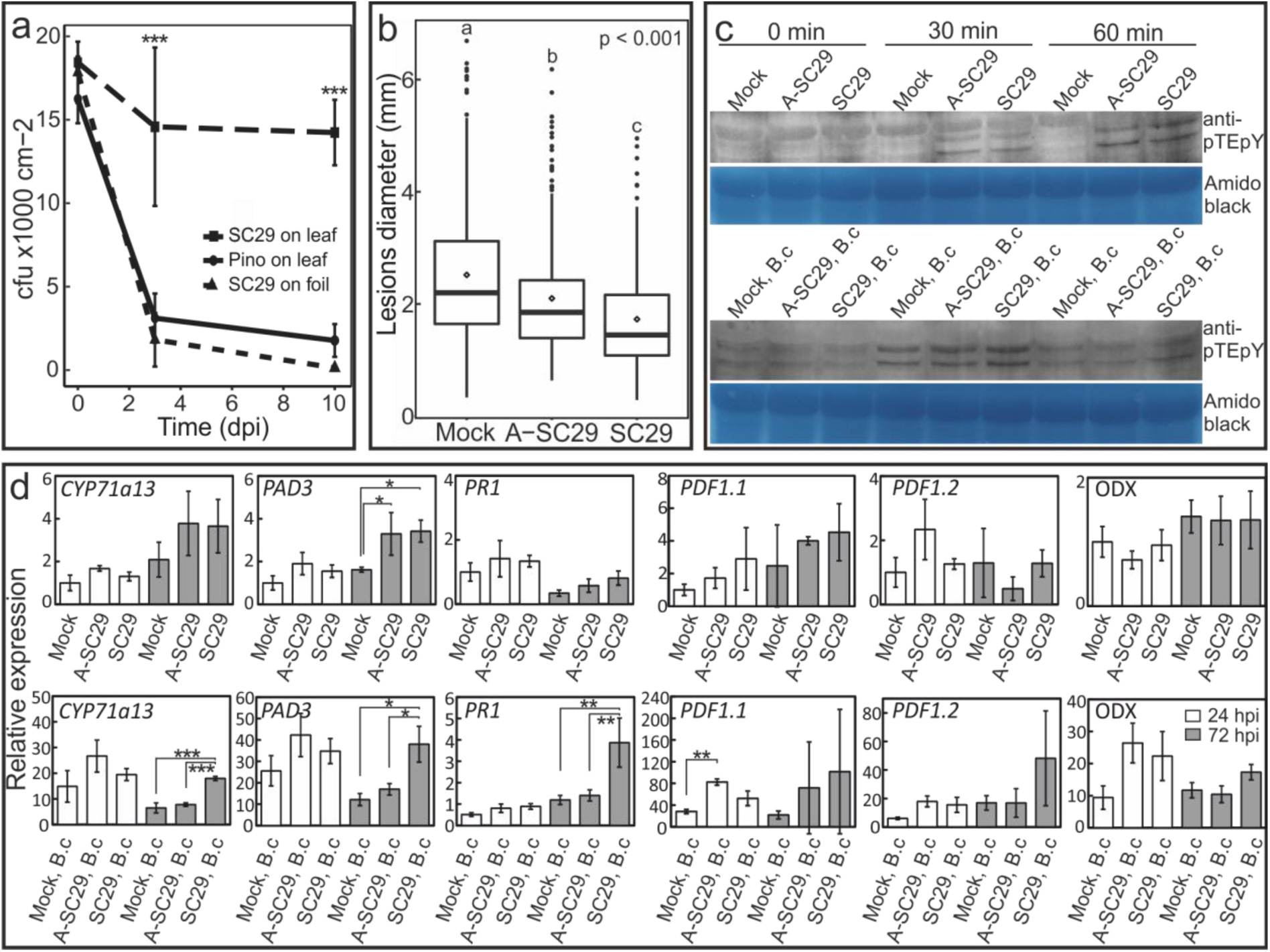
SC29 interaction with *Arabidopsis* immune signalling. a) Persistence assay of SC29 and *P. inouyei* (Pino) on the surface of *Arabidopsis* leaves aseptically grown on 0.5xMS agar. Cells were re-isolated from drop inoculated plants at the indicated times and plated to determine cell numbers, which are presented as the number of colony forming units (CFUx1000 cm^-2^). Persistence on sterile foil was used as a control for the ability to survive non-specifically on surfaces. Pooled data from five independent biological repeats (n=25 total) were analysed by computing a linear mixed model in R (3.5.1). b) Lesion diameters of *Botrytis* drop infections on *Arabidopsis* leaves pre-treated with water (mock), autoclave killed SC29 (A-SC29), or live SC29 cells. The bars represent the mean lesion size ±SD (n=24 total). Statistics performed with pooled data from six independent biological repeats by computing a linear mixed model in R. c) *Arabidopsis* MAPK3/6 activation was monitored by western blot with anti-phospho (activated) MAPK antibodies following pre-treatment with A-SC29 or live SC29 at 30 and 60 min. Equal loading was confirmed by amido black staining. Three independent experiments were repeated with the same result, representative results are shown. d) Relative gene expression of plant defence signalling genes 24 (white bars) and 72 (grey bars) hours post infection (hpi) following SC29 treatments only (top row) and after *Botrytis* infections with and without SC29 pre-treatments (bottom row). Gene expression was normalized to water control 24 hpi for each gene. Data are presented as mean relative expression levels ±SD of three pooled biological repeats. Statistics was performed with glht package in R (3.5.1) with fdr method for adjusted p-values. Key to abbreviations: A-SC29, autoclaved SC29; B.c, *Botrytis cinerea.* * p < 0.05; ** p < 0.01; *** p < 0.001.

### SC29 activates MAPK and hormone immune signalling pathways

To define the SC29-*Arabidopsis* interaction we tested the ability of SC29 to enhance resistance against a broad-host-range fungal pathogen. Plants were pre-inoculated with water, SC29, or SC29 killed by autoclaving (A-SC29), prior to infection with the necrotrophic pathogen *Botrytis cinerea* and disease progression was monitored as the size of *Botrytis*-induced spreading lesions. Significantly smaller lesion diameters in *Arabidopsis* pre-treated with both live and killed SC29 (Fig. 2b) were observed, suggesting increased resistance. Additionally, live SC29 pre-treated plants had significantly smaller lesions than those treated with A-SC29. This suggests that yeast MAMP molecules liberated from dead SC29 could trigger plant immunity, but the full effect required the live SC29. Co-cultivation on artificial media demonstrated that SC29 was not able to inhibit *B. cinerea* growth (Fig. S5), ruling out possible direct interactions. These results suggest SC29 interacts with *Arabidopsis* to activate immune signalling.

To test early immune signalling, we assayed MAPK activation in response to SC29 pre-treatment, both alone and with subsequent *Botrytis* infection. Western blots probed with an antibody specific for the active phosphorylation site of MAPKs revealed activation of *Arabidopsis* MAPK3 and MAPK6 (MAPK3/6) in response to treatment by both live and dead SC29 at 60 min post infection (Fig. 2c). *Botrytis* infection also resulted in MAPK3/6 activation.

To further explore activation of defence signalling, we employed qPCR to monitor expression of defence signalling marker genes (for gene names and AGI codes, see File S2). Strikingly, we observed evidence of immune priming, seen as enhanced *Botrytis*-induced transcriptional responses, in samples pre-treated with both live and dead SC29 (Fig. 2d), for *PDF1.1*, a jasmonic acid (JA) marker; *CYP71a13* and *PAD3*, markers of camalexin - the primary antimicrobial compound in *Arabidopsis*; and *PR1*, a marker of salicylic acid (SA). Live and dead SC29 alone had limited effect, showing induction at 72 hpi of *PAD3* (Fig. 2d). These results indicate SC29 was able to activate some of the known canonical defense pathways and implicates JA, SA, and camalexin in the *Arabidopsis* response to this yeast. Markers of other signaling pathways such as ethylene, ABA, auxin, and cytokinin exhibited no significant changes (Fig. S6), suggesting the SC29 response is independent of these pathways.

### Gene expansion and loss

We identified candidate effector-like proteins (CELPs) in the genomes of *Protomyces* spp. and categorized them as small secreted proteins (SSPs), some of which were cysteine-rich (CSSPs; Table S2). All *Protomyces* had a higher number of SSPs and CSSPs compared to *Taphrina* spp. and the non-pathogenic *Schizosaccharomyces* spp., exceptionally, *T. flavorubra* had more SSPs than Pgra (Table S2). All *Protomyces* SSPs had similar features and were slightly smaller and more cysteine-rich compared to *T. deformans* (Fig. S7). Positive hits in the PHIbase indicated the presence of conserved fungal virulence proteins or effector-like proteins in *Protomyces* genomes (File S5). Identification of conserved protein domains in *Protomyces* CELPs (File S1) revealed the remarkable lack of LysM domains. However, the carbohydrate-binding, legume-like lectin (PF03388.13) domain was present in *Protomyces*, except Ppac. Topology of the phylogenetic tree constructed with SSPs (File S6), suggests that Pmac, Pinu, and Pgra CELP evolution diverged early from the other *Protomyces* species.

BadiRate analysis for estimation of gene family turnover rates (28) with SSP gene families (orthogroups, OGs) identified eight significant expansions (File S6), including one species-specific expansion in SC29 in SSP OG12. Public database queries with genes in the expanded OGs revealed that they are all *Protomyces*-specific genes with no conserved domains. These results suggest that although Pgra, Pino, Plac, and Ppac are all adapted to Compositae family hosts, CELPs in Pgra have a distinct evolutionary history. The expansion in SC29 suggests potential adaption to a different lifestyle or host, prompting us to seek additional evidence of this.

Gene loss is a feature in fungal genomes that have undergone a host jump (45). To test for this, gene content was compared between the closely related SC29, Pino, Plac, and Ppac (Fig. 3). Altogether 283 SSP OGs were absent from SC29 but present in the other three species, while 24, 40, and 22 were missing in Pino, Ppac, and Plac, respectively, but present in all others (Fig. 3a). Many, but not all, of these SSPs were cysteine rich (Fig. 3b). Although loss of a small number of genes from each OG were not detected by BadiRate as significant, cumulatively these losses are likely biologically relevant.

**Figure 3.**
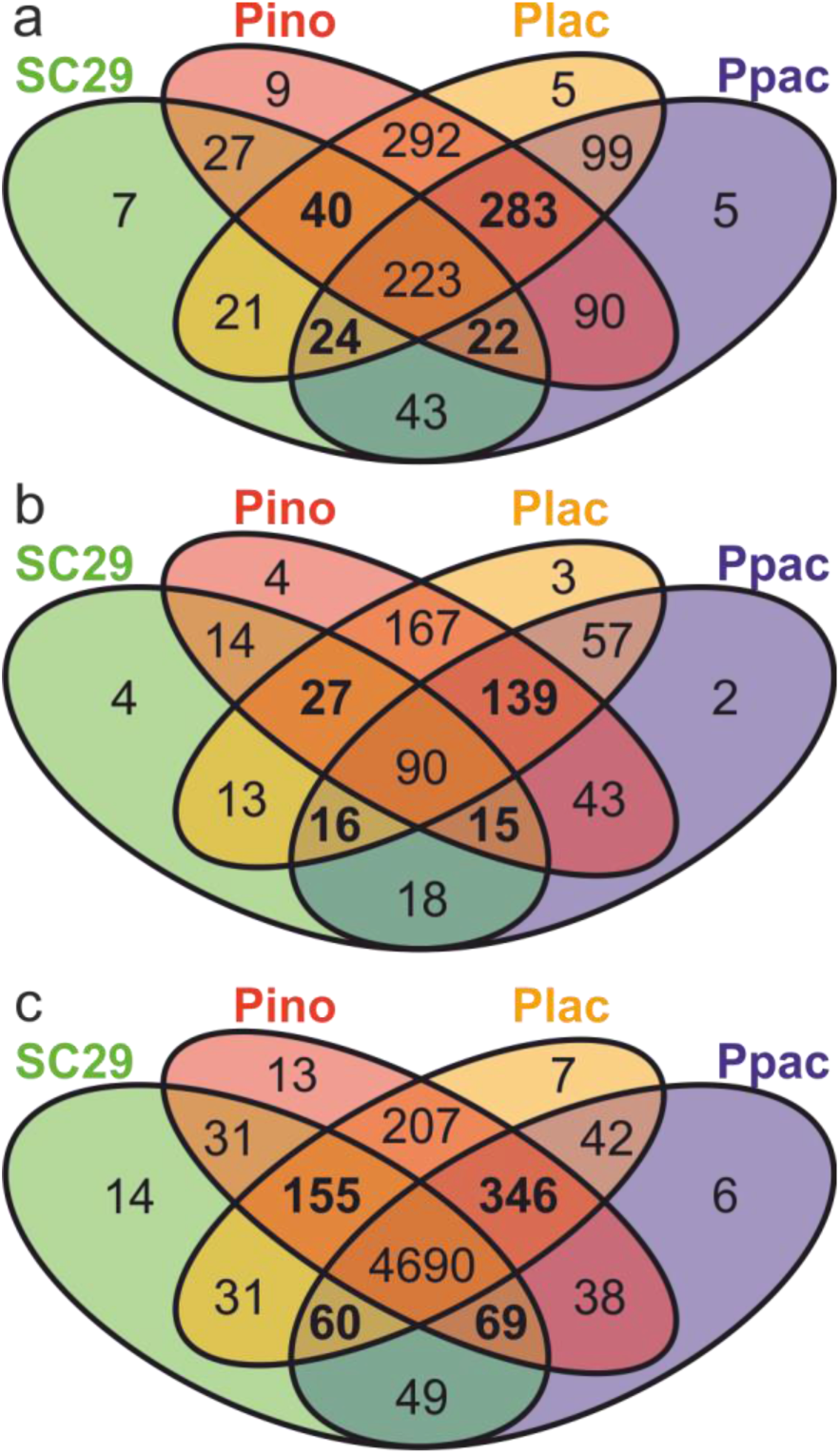
Shared orthologous gene clusters among selected *Protomyces* spp. The species used form a closely related clade and include; *Protomyces* strain C29 (SC29), *P. inouyei* (Pino), *P. lactucaedebilis* (Plac), and *P. pachydermus* (Ppac). a) Orthologous clusters of candidate effector-like proteins (CELPs) of the small secreted protein (SSP) class. b) Orthologous clusters of CELPs of the cysteine-rich SSPs (CSSPs) class. This is a subset of the genes in (a). c) Orthologous clusters found in all annotated genes in the genomes of these species. Clusters common to three of the four species are indicated in bold.

Gene loss in SC29 was also evident at the whole genome level. SC29 only encoded 5514 annotated genes, compared to 5956, 5612, and 5952 for Pino, Ppac, and Plac, respectively (Table 1). Comparisons of the whole genome data revealed 346 OGs missing from SC29, while 60,155, and 69 were absent from for Pino, Ppac, and Plac, respectively (Fig. 3c). Gene ontology (GO) analysis of genes absent from SC29 indicated enrichment in GO:0036267 (invasive filamentous growth; File S7). These four genomes show high levels of similarity as evaluated by ANI, AAI (Table 1), and synteny (Fig. S1). Their genome completeness levels estimated with BUSCO were similar (86.0-86.3%), indicating similar amounts of unannotated genes.

Genes found only in one of these four species were also identified; 14 OGs (32 genes), 13 OGs (28 genes), six OGs (16 genes), and seven OGs (19 genes), were found to be species specific for SC29, Pino, Ppac, and Plac, respectively (Fig. 3c). SC29 specific genes were enriched for GO:0036349 (galactose-specific flocculation) and GO:1900233 (regulation of biofilm formation on inanimate substrates) (File S7).

Taken together, we conclude that SC29 exhibits patters of gene loss. These genomic changes are consistent with an adaptation to a new host and new lifestyle.

### Carotenoids and auxin-like compounds in Protomyces

Assays of *Protomyces* indolic-compound production showed variation among species, ranging from 0.9 to 7.7 µg/ml when cultured in GYP medium and from 2.3 to 25.6 µg/ml with added tryptophan (Fig. 4a). Indolic-compound production was elevated in all cultures with additional tryptophan. *Protomyces*-derived indolic compounds in culture supernatants activated the artificial auxin-responsive promoter (*DR5*) in transgenic *Arabidopsis* bearing a *DR5∷GUS* fusion, resulting in deposition of blue GUS stain (Fig. 4b). Additionally, we assayed auxin activity phenotypically, as the induction of root hair growth by *Protomyces* culture filtrates (Fig. 4c). Taken together, these results indicate that *Protomyces* spp. produce indolic compounds with auxin-like activity, prompting us to identify the possible indole acetic acid (IAA) biosynthesis pathways encoded in these genomes. Of the five queried pathways, genes encoding all required enzymes were present only for the indole-3-pyruvic acid (IPyA) pathway (Table S3).

**Figure 4.**
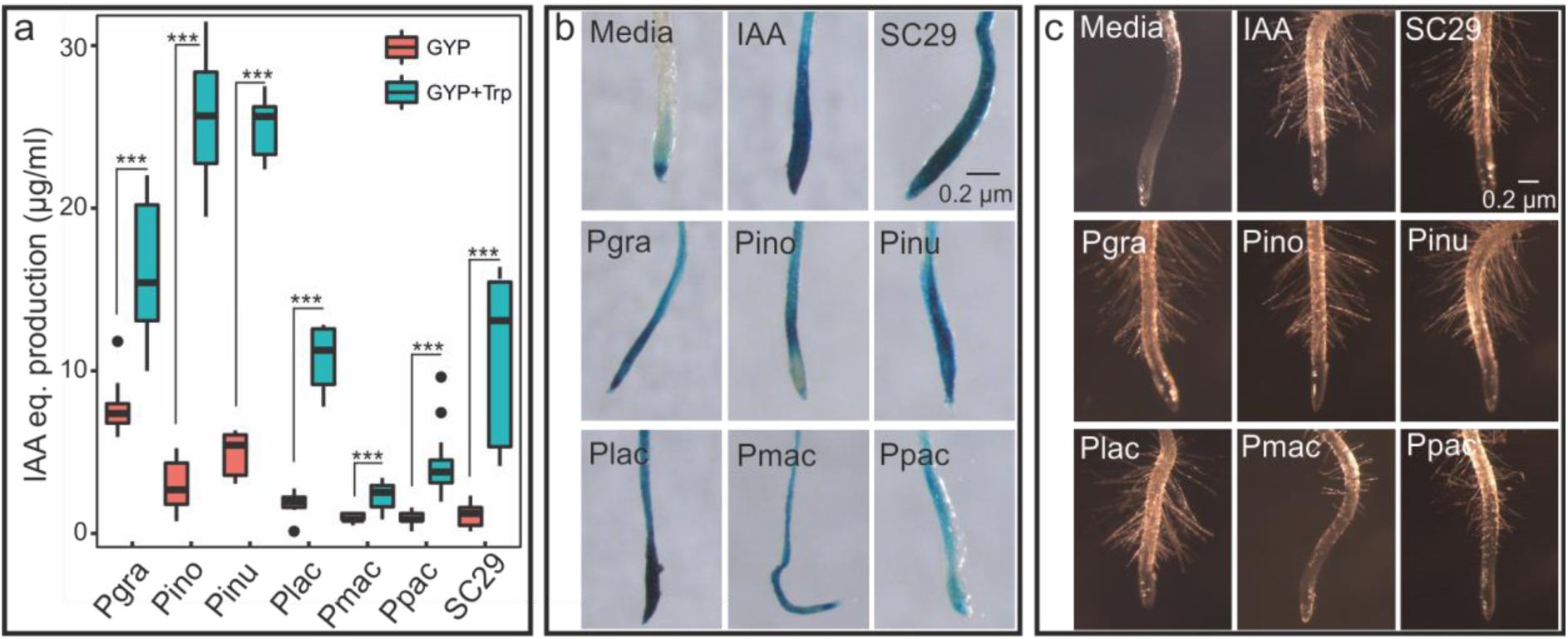
The production of auxin-like compounds by *Protomyces*. Species used and their abbreviations are: *Protomyces* strain C29 isolated from wild arabidopsis (SC29), *P. gravidus* (Pgra), *P. inouyei* (Pino), *P. inundatus* (Pinu), *P. lactucaedebilis* (Plac), *P. macrosporus* (Pmac) and *P. pachydermus* (Ppac). a) Quantification of indolic compounds produced by each *Protomyces* species using the Salkowski reagent assay on supernatants from five-day-old cultures in the specified liquid media [glucose yeast extract peptone (GYP) and GYP with additional tryptophan (GYP+Trp)]. Results are presented as equivalents (IAA eq.) from a standard curve with the auxin, indole acetic acid (IAA) and are means ±SD. Statistics were performed with t-test in R (3.5.1), *** p < 0.001. b) *In vivo* activation of the auxin transcriptional response is shown in roots of *Arabidopsis* bearing an artificial auxin-inducible promoter-reporter fusion (*DR5:GUS*) treated with sterile filtered culture supernatants. Auxin-induced promoter activity is visualized by deposition of blue stain indicating the presence of β-glucuronidase (GUS) reporter activity. c) Root hair formation as a biological assay for auxin activity. Experimental details are as in b. Uncultured media was used as a negative control (media) and IAA (5µM in b and 1µM in c) was used as a positive control (IAA). Three independent biological repeats were conducted for all experiments.

We further queried for possible carotenoid biosynthesis pathways within these genomes (Fig. S8). These were found in all seven *Protomyces* spp. with homologs of key enzymes phytoene desaturase (albino-1) and lycopene beta-cyclase (albino-2), but not lycopene epsilon-cyclase (CrtL-e).

### Other features of Protomyces genomes

To probe for genomic features associated with the lifestyle of *Protomyces* spp., BadiRate was utilized in comparisons within and between these *Protomyces* species and non-pathogenic ascomycete yeasts using proteome data (File S6). Selected gene families with statistically significant lineage-specific expansions related to *Protomyces* lifestyle are highlighted below.

Orthogroup (OG)184 are a group I sensory histidine kinases. OG184 is absent from *Schizosaccharomyces* spp., present in variable copy number in all *Taphrina* and *Protomyces;* in SC29 it shows a significant expansion to five copies (Table S4). We manually curated all hybrid histidine kinases (HHKs) present in these *Protomyces* genomes (Table S4; File S8). Remarkably, in Pmac a dual HHK (g3206) was found immediately adjacent to another type XI HHK (g3205). Several adjacent HHK gene pairs were found in other *Protomyces;* in Pinu two type XI HHKs (g5765-g5766) and two pairs in SC29 involving the type I HHK gene family (the expanded OG184) adjacent to a type IX and XI HHKs (g3356-g3357 and g3889-g3890, respectively).

OG3409 and OG226 encode two related pectin degradation activities. OG102 encodes a family of secreted subtilisin-like serine proteases. These OGs both are absent from *Taphrina* and *Schizosaccharomyces* species, present in all *Protomyces* species, and show significant expansions in some *Protomyces* lineages (File S6). OG46 encodes the serine protease component of the SPS sensor. Members of this gene family are present in a single copy in two *Taphrina* species and show a significant expansion in all but one *Protomyces*, where they are present in high (three to nine) copy number. OG53 encodes cutinase gene palindrome-binding proteins, which were present with one or two copies in all species examined and significantly expanded to seven copies in SC29.

Finally, we investigated carbohydrate-active enzymes (CAZymes) in *Protomyces* genomes using dbCAN (41). All *Protomyces* genomes harboured a similar and relatively low number of CAZymes (241-261 total; File S9).

## DISCUSSION

### Phylogenetic implications

Our results have several implications for the phylogenetics of the genus *Protomyces*, which have been handled elsewhere (12).

### An Arabidopsis associated Protomyces

We previously presented four *Protomyces* strains isolated from *Arabidopsis* at two distinct sites in Helsinki, Finland (13). Here we expand those results with *Arabidopsis*-associated *Protomyces* evidence from ITS sequencing in Germany (Fig. S3). The occurrence of *Protomyces* on *Arabidopsis*, together with the ability of SC29 to both persist in the *Arabidopsis* phylloplane and interact with the *Arabidopsis* immune system, give evidence that this is a true interaction. Although, at this time, we cannot claim that *Arabidopsis* is the only or the primary host. Still, this is the first instance of a *Protomyces* species associated with a host plant outside of the Umbelliferae or Compositae families and suggests this novel *Protomyces* sp. may have become adapted to a new host or hosts. The novel host and genomic signatures observed in its genome prompt us to propose the hypothesis that SC29 may have undergone a host jump.

In defining this interaction, we have established a model experimental system with a plant-associated yeast and the genetic model-plant *Arabidopsis*, which will facilitate future genomic studies into the biology and lifestyles of plant associated species in the Taphrinomycotina, the evolution of fungal virulence, host interactions with phyllosphere yeasts, and plant immunity against yeasts. The activation of *Arabidopsis* defence signalling pathways by SC29 provides evidence of yeast-specific MAMPs that can trigger plant immunity.

Remarkably, human pathogenic yeasts lack LysM effectors (46). The human innate immune system deploys a distinct set of pattern recognition receptors (PRRs) activated by mostly mannose containing linkages that are highly abundant in the outer layers of cell walls of pathogenic yeasts (47, 48). Such plant PRRs responsible for detecting yeast MAMPs remain unknown, although it has been demonstrated that plants can detect the presence of yeasts and activate immune signalling pathways. Engagement of *Arabidopsis* immune signalling has been previously observed with autoclaved cell suspensions of *S. cerevisiae* resulting in the activation of SA signalling, camalexin biosynthesis, and enhanced resistance against other pathogens (49). Accordingly, with SC29 we observed activation of SA and camalexin pathways and enhanced *Botrytis* resistance.

*Protomyces* spp. invade their hosts in their hyphal form. The genomes of filamentous (hyphal) fungal pathogens have chitin containing cell walls and frequently contain lysine motif (LysM) domain effectors involved in sequestering chitin or otherwise blocking immune responses elicited by this MAMP (46). The absence of LysM domain containing CELPs in *Protomyces* spp. suggests that chitin is of lesser importance for their host interactions. Indeed, CELPs bearing another conserved carbohydrate-binding domain, the legume (L)-type lectin domain were present in all but one *Protomyces* genome. This lectin domain mediates the binding of mannose linkages (50-52). Pinu has been shown to have a cell wall composed of glucan and mannose, similar to other yeasts (53) and *S. pombe* cells has been shown to have very little chitin, which is found restricted only to certain structures (54). Taken together, these results further support the importance of mannose-linkages, or other yeast MAMPs, in the interactions of *Protomyces* with their plant hosts. Forward and reverse genetic screens aimed at discovering PRRs involved in detecting yeast MAMPs (yeast cell wall components) are underway in our laboratory.

### Comparative genomics within the Protomyces

The phyllosphere is estimated to be one of the largest microbial habitats on earth with an estimated size of 10^9^ km^2^ (55). However, this habitat presents abiotic hazards such as lack of nutrients, exposure to full solar irradiation, extreme temperature fluctuations, and long periods of water shortage punctuated by periodic deluge of raindrops, which threaten to dislodge microbes (55, 56). Biotic threats are also present; for instance, microbe-microbe interactions such as competition for resource acquisition (57) and growth inhibition by antibiotics (58). Although not yet documented in the phylloplane, effector proteins can be deployed in direct inter-microbial competition (59-61). The host is also a threat; host derived antimicrobial secondary metabolites are present in the phyllosphere (62) and increasingly a picture is emerging where changes in the cuticle can trigger plant immune signalling, extending plant immune surveillance out into the phyllosphere (63-65). Thus, the phyllosphere is a remarkably hostile environment that presents a significant barrier to be overcome by resident microorganisms and pathogens utilizing this space for host access. Accordingly, phyllosphere microbiome communities comprise microbes specifically adapted to this environment (10, 55).

Genome sequencing enables the elucidation of genomic features of plant-associated microbes. Significant advances have been made, especially in root associated microbial communities, especially prokaryotes (66, 67). Phyllosphere microbes, including yeasts, are now gaining considerable interest, but our understanding still lags behind. Also, information on the role of phylloplane residency as a yeast in the *Protomyces* lifecycle is not available (2, 3). Our genome data presents an opportunity to address these question by probing the genomes of *Protomyces* species for evidence of phyllosphere adaptations.

Several features related to nutrient acquisition were found in these genomes. Gene families belonging to the SPS sensor were expanded in these *Protomyces* spp. In *Candida albicans* the SPS system has also been implicated in immune evasion and nutrient acquisition, by sensing extracellular amino acids (68). Other expanded gene families include: pectinase gene families that may be involved in utilization of pectin as a nutrient in the plant phylloplane and a family of secreted subtilisin-like serine proteases previously linked with nutrient acquisition during soil-residency (69). Finally, carbohydrate-active enzyme (CAZyme) gene content was defined; all *Protomyces* species had similar and low number of CAZymes relative to other fungal pathogens, consistent with their small genomes and facultative-biotrophic pathogen lifestyles (70).

Auxin of microbial origin is multifunctional controlling growth promotion by beneficial microbes (71, 72) and immune suppression by pathogens (73-75). Auxins in the phyllosphere can inhibit bacteria (76) and enhance host nutrient leaching via cell wall loosening and sugar release (55, 77, 78). Our data suggests that *Protomyces* produce IAA. The pathways responsible for fungal IAA biosynthesis are not well defined; several pathways are suggested (79, 80). Exogenous tryptophan feeding experiments suggest a tryptophan-dependent IAA biosynthesis pathway in *Protomyces* spp. Accordingly, these genomes all encoded a complete indole-3-pyruvic acid (IPyA) pathway (Table S3), suggesting this as a candidate IAA biosynthesis pathway in *Protomyces* spp. The IPyA pathway has been suggested for other Taphrinomycotina species, *T. deformans* (5, 8) and three other *Taphrina* species (8), based on the presence of orthologs of TAM and IAD genes, similar to the pathways previously found in the Basidiomycete, *U. maydis* (81). Tsai *et al* also found orthologs of the YUC Flavin monoxygenase in four *Taphrina* species, suggesting the presence of an additional plant-like IAA biosynthesis pathway (8); however, YUC orthologs were absent from *Protomyces* spp. genomes (Table S3).

Carotenoids are important stress protectants in phylloplane yeasts (82, 83) and are responsible for coloured colonies observed in *Protomyces* cultures (1). Three *Protomyces* spp. (Pino, Ppac, and Plac) previously assayed produce high levels (65-99 ug/g dry weight) of carotenoids (84). Carotenoid biosynthesis pathways present in *Protomyces* spp. suggest that all produce lycopene, γ-carotene, and β-carotene, but not δ-carotene or α-carotene.

Some phylloplane adaptations were found that were specific to SC29, consistent with its demonstrated ability to persist on the leaf surface of *Arabidopsis*. One expanded family of genes encoded the DNA repair protein Mre11, which is the nuclease subunit of the widely conserved double-strand break repair MRX complex with Rad50p and Xrs2p (85). DNA repair represents a phyllosphere protective mechanism against damage from solar irradiation and reactive oxygen species produced by plant defence responses (55). The cutinase transcription factor 1 (CTF1) gene family was specifically expanded in SC29. CTF1 is required for induction of fungal cutinase by plant hydroxyl fatty acids from cutin (86, 87), a component of the plant cuticle and potential energy source.

### Sensory histidine kinases

HHKs are important sensory proteins transmitting both internal and environmental information in the cell (40, 88). Most HHKs groups were found in *Protomyces* spp. genomes, with some exceptions (Table S6). Several genomic features suggest that the HHKs are actively evolving in *Protomyces* spp. Groups III and X are involved in regulation of pathogenicity and morphogenesis in pathogenic fungi (40). The usual single copy of group III HHKs normally in pathogenic fungi (40), is doubled in several *Protomyces* spp. Group I HHK genes have undergone a significantly expansion in SC29. Their function is not well studied (40); but may be related to development of virulence (89). Two of these group I HHK genes are immediately adjacent to other HHKs in SC29. Tandemly duplicated type XI HKKs were also observed in Pinu. Remarkably, in Pmac a dual HKK was adjacent to a type XI HKK. Dual HHKs have only been found in the genomes of fungi in the Basidiomycota. *Protomyces* belong to the Taphrinomycotina, a subphylum of the Ascomycota noted for unique species with ancestral characteristics. This first instance of a dual HHK in an ascomycete is consistent with the observation of basidiomycete-like and ancestral features in other fungi in the order Taphrinales (4, 90). *Protomyces* spp. also possess other basidiomycete characteristics. The genomes of all *Protomyces* spp. have a GC content over 50% (Table 1), which is more consistent with basidiomycetes than ascomycetes (91). Both *Protomyces* and *Taphrina* spp. exhibit an enteroblastic budding, a basidiomycete characteristic, in contrast to holoblastic budding in ascomycetes (91, 92). The naked asci of these species has been described as basidiospore-like (93, 94). Moreover, Q-10 is the major ubiquinone system in *Protomyces* and *Taphrina* spp. (91, 95). Finally both possess resting cells or reproductive structures that are described as thick walled “chlamydospores” (96, 97).

Two GO categories were enriched in the SC29 specific genes, galactose-specific flocculation and positive regulation of single-species biofilm formation on inanimate substrate. These SC29 specific genes are related to the *S. pombe* galactose-specific cell agglutination protein (gsf2), an outer PM protein promoting cell-to-cell adhesion (98, 99) and the *Candida albicans* biofilm regulator 1 (BRG1) transcription factor (100, 101). Microbial aggregates and biofilms facilitate water retention and wetting in the phylloplane; in addition to their roles in attachment these are adaptations promoting drought tolerance and nutrient leaching (55).

Taken together, our results support that SC29 is able to survive in the *Arabidopsis* phyllosphere. This is similar to all known *Taphrina* species where phylloplane residency feature prominently in the lifecycle. SC29 may be analogous to several known *Taphrina* species originally assigned to the defunct genus *Lalaria*, which are *Taphrina* isolated in the yeast (anamorphic) state from asymptomatic hosts (102, 103). These *Taphrina* not associated with host disease are thought to have either lost virulence and live only in the phylloplane or to be pathogenic on unknown hosts and utilize phylloplane residency on presumed alternate hosts (4, 103). Given the relatedness of *Taphrina* and *Protomyces* and their similarity in lifestyle and pathogenicity strategy, the discovery of phyllosphere resident *Protomyces* species is plausible. Although the anamorph genus *Lalaria* has been rendered invalid by abandonment of the dual name system, the questions raised by the phylloplane-resident lifestyle that defined the *Lalaria* remain relevant to understanding the biology of yeasts of the order Taphrinales.

SC29 is the first *Protomyces* found to be able to persist in the phyllosphere of its presumed host, or alternate host. This together with phyllosphere genomic adaptations found in the genomes of all *Protomyces* species illustrates the importance of phylloplane survival for the *Protomyces* lifecycle, for instance in the pre-infection period while waiting for favourable infection conditions or compatible strains for conjugation. This is consistent with the ability of species the genera *Protomyces* and *Taphrina* to forcefully eject their spores from ascogenous cells (2), supporting that *Protomyces* species, like *Taphrina* species, are wind dispersed and colonize their hosts via the phylloplane.

## ACKNOWLEDGMENTS

We thank Tuomas Puukko, Airi Lamminmäki, and Leena Grönholm, for excellent technical support and Katariina Vuorinen for assistance with infection assays, Mikeal Broché and Tiina Blomster for advice of qPCR, Julia Krasensky-Wrzaczek for instruction on MAPKs work, Ansa Palojärvi (Joint ‘SUCCESS’ project) for the control *Paenibacillus* strain with antifungal activity, and the personnel of the DNA sequencing and genomics laboratory performing NGS sequencing. This work was supported by the following grants: Academy of Finland Fellowship (decisions no. 251397, 256073 and 283254) to KO and the Academy of Finland Center of Excellence in Primary Producers 2014-2019 (decisions #271832 and 307335). KW, SR, and OM are members of the University of Helsinki Doctoral Programs in Plant Science (DPPS) and AA in Microbiology and Biotechnology (MBDP). Computing resources provided by the Finnish IT Center for Science (CSC; www.csc.fi) are gratefully acknowledged. We wish to thank Prof. Dr. Dominik Begerow and Prof. Daniel Croll for their critical comments on this manuscript during the external examination of Kai Wang’s PhD thesis.

