## Supplemental Figures and Tables for "A novel phyllosphere resident *Protomyces* species that interacts with the *Arabidopsis* immune system"

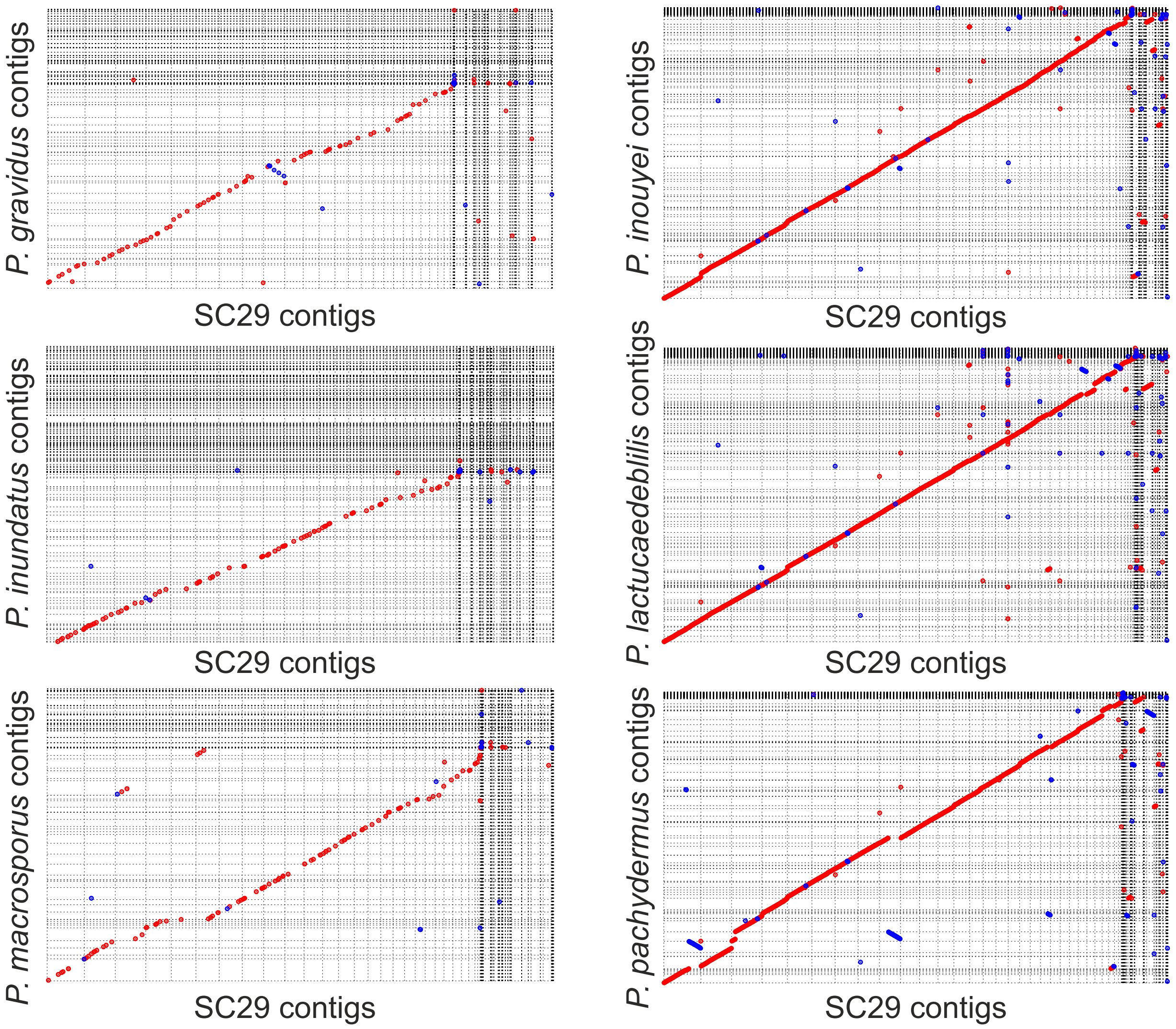


**Figure S1.** Dot-plots of whole genome alignments between SC29 and six other *Protomyces* species. Mummer 4.0.0beta2 (nucmer) was used for genome assembly comparison with default settings. Forward alignment is plotted in red and reverse alignment in blue. Optimal co-linear order of contigs was shaped with mummerplot (parameter --fat). Mummerplot output .ps files were viewed and labeled with CorelDRAW2018.

a

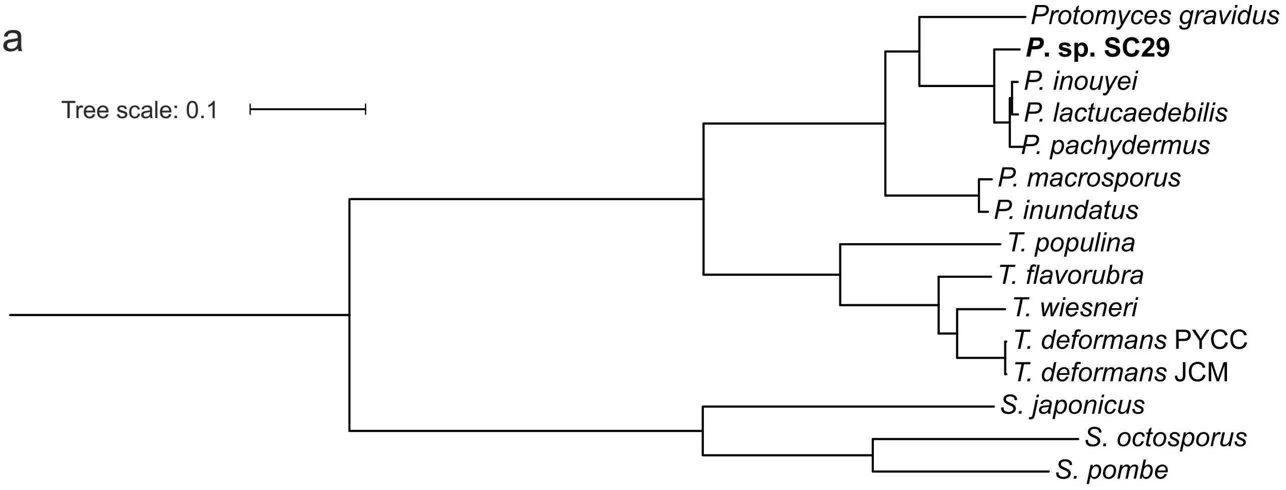


b

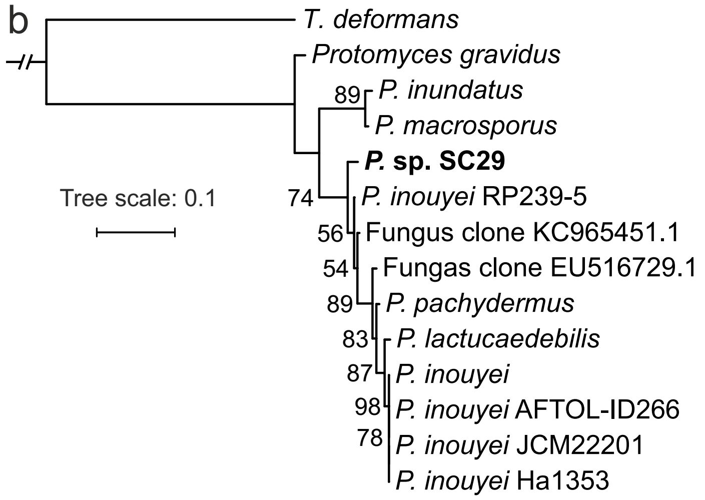


c

| Species | *No. of isolate hits |
|---|---|
| *P.* sp. SC29 | 6 |
| *P. gravidus* | 0 |
| *P. inoyei* | 2 |
| *P. inudatus* | 0 |
| *P. lactucaedebilis* | 2 |
| *P. macrosporus* | 0 |
| *P. pachydermus* | 1 |

**Figure S2.** Phylogenies and *Protomyces* spp. in environmental sequencing data. a) Genome-wide phylogenetic tree of the genera *Protomyces* and *Taphrina*, built with the Bayesian inference method with Mrbayes 3.2.7a. General time reversible (GTR) model and invgamma were chosen. Two independent analyses were started simultaneously with 10000 generations were set for analysis run. The output .con.tre files were viewed with iTol and edited with Coreldraw2018. b) ITS phylogenetic tree constructed with the maximum-likelihood method (RAxML) using *Protomyces* reference spp., SC29, and six SC29-like *Protomyces* isolates from environmental samples. c), Numbers of hits for each *Protomyces* species from searches of environmental ITS metagenomic library sequencing and environmental isolates data queried at NCBI. The criteria for positive hits here are ITS similarity >97% and coverage >95%. Only hits to environmental strains are considered in this list. The NCBI accession numbers for the isolates where: for SC29 - LT602859.1, MF615097.1, MK045398.1, KC965451.1, MF615002.1, KX067824.1; for *P. inouyei* - KU134840.1, EU516729.1; for *P. inundatus* - MK937059.1; for *P. lactucaedebilis*, KU134840.1, EU516729.1; for P. pachydermus - EU516729.1.

**Figure S3.** Occurrence of *Protomyces* OTUs over time in *Arabidopsis thaliana* common garden field planting experiments in Cologne, Germany. a), Experiments were run during the *Arabidopsis* growing season (November to March) and were repeated three times in the years 2014-2016. Pooled data from three years indicate the abundance of *Protomyces* OTUs determined by ITS barcode library sequencing. *Arabidopsis* accesssions used were Col-0 (C), Ksk1 (K), Sf-1 (S), Ws-0 (W). Letters above the plots for each month indicate significance groups (p<0.05) by one-way ANOVA + Tukey HSD. b), An ITS1 based phylogenetic tree with *Protomyces* OTUs sequenced in Cologne, SC29, and six reference *Protomyces* species. The phylogeny was built with neighbor-joining method in ClustalX2.1 using *Protomyces gravidus* as an outgroup. The position of *Protomyces* sp. SC29 was highlighted in bold.

a

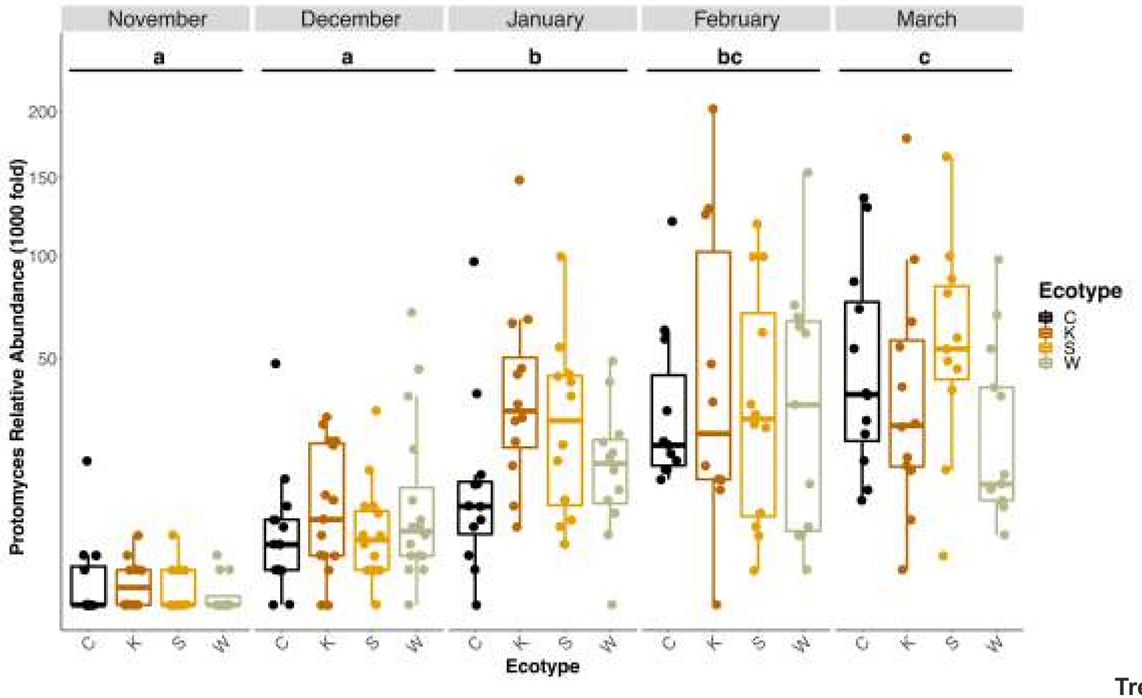


b

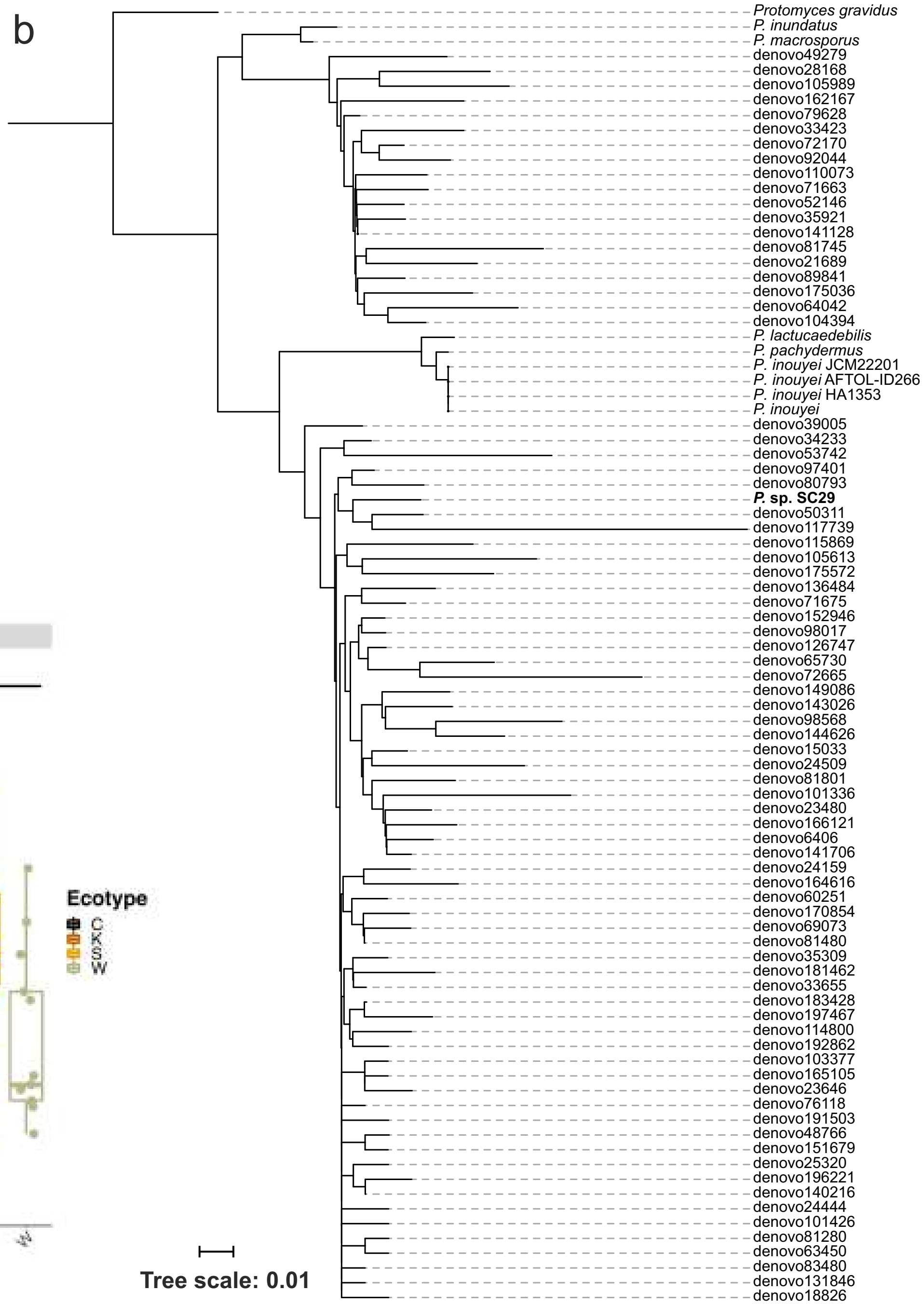

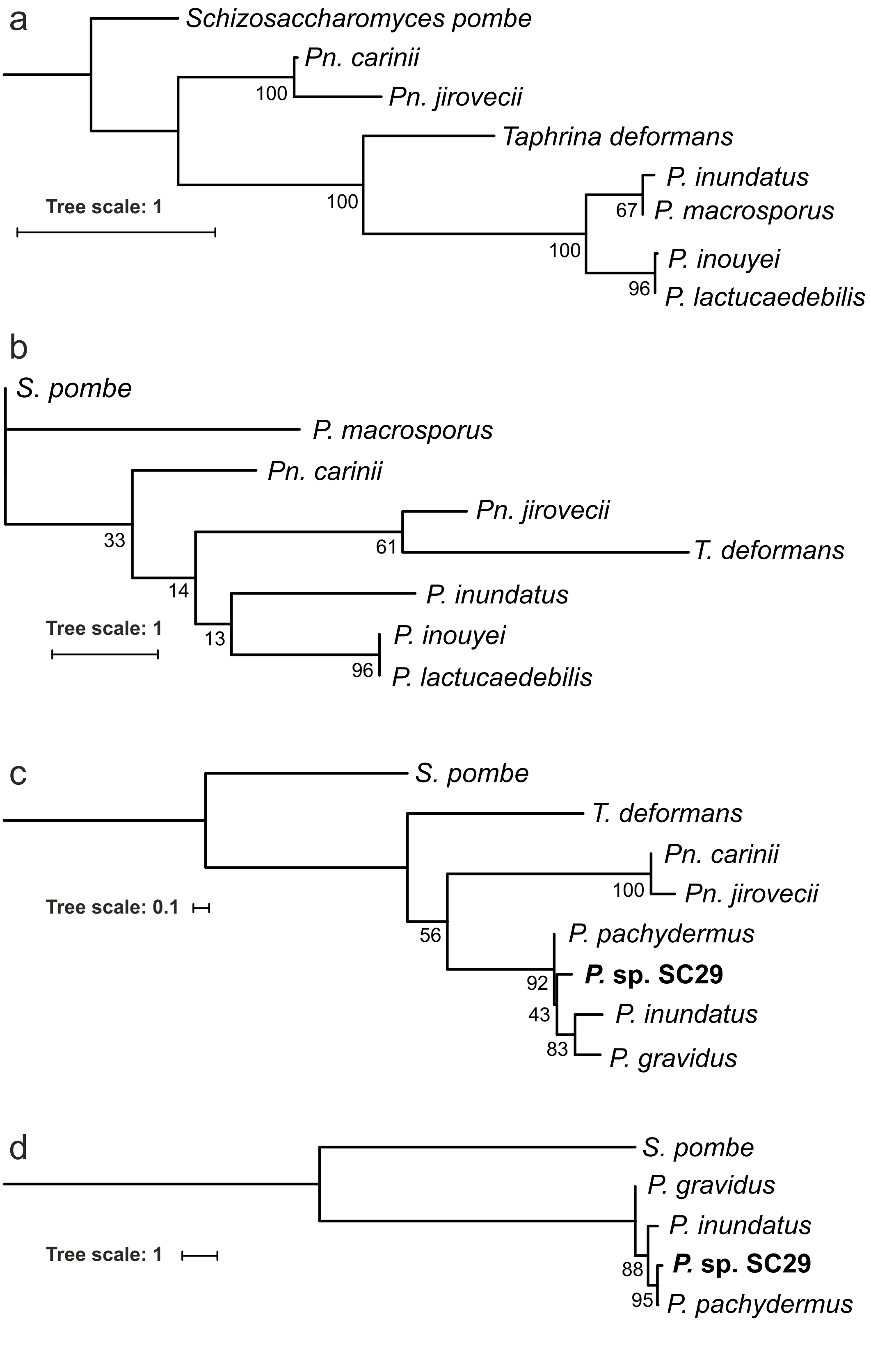
a
Schizosaccharomyces pombe
Pn. carinii
100
Pn. jirovecii
Taphrina deformans
P. inundatus
Tree scale: 1
100
67
P. macrosporus
100
P. inouyei
96
P. lactucaedebilis
b
S. pombe
P. macrosporus
Pn. carinii
Pn. jirovecii
33
61
T. deformans
14
P. inundatus
Tree scale: 1
13
P. inouyei
96
P. lactucaedebilis
c
S. pombe
T. deformans
Pn. carinii
100
Pn. jirovecii
Tree scale: 0.1
56
P. pachydermus
92
P. sp. SC29
43
P. inundatus
83
P. gravidus
d
S. pombe
P. gravidus
P. inundatus
Tree scale: 1
88
P. sp. SC29
95
P. pachydermus

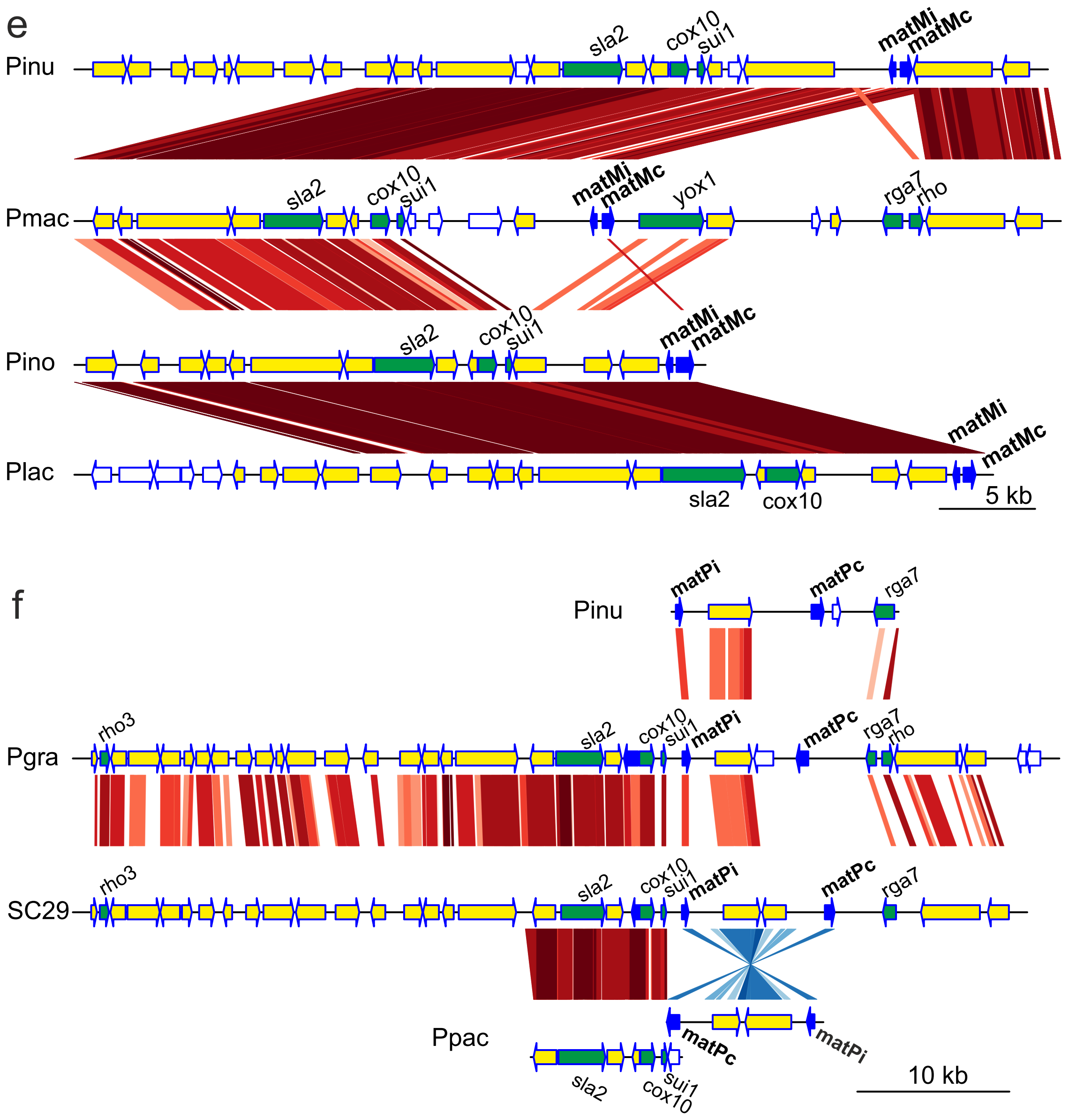


**Figure S4.** Mating (*mat*) genes and MAT loci of *Protomyces* spp. Mating genes of *Protomyces* spp. were identified based on similarity to known mating genes and the presence of adjacent MAT loci associated genes such *rga7*, *sla2*, *cox10*, and *sui1*. Mating genes from *Taphrina deformans*, *Schizosaccharomyces pombe*, and *Pneumocystis* spp. were used as queries. Maximum likelihood phylogenetic trees using protein sequences of *Protomyces* spp. encoded by the mating genes are shown for *matMc* (a), *matMi* (b), *matPi* (c), *matPc* (d). *Schizosaccharomyces pombe* was used as an outgroup. Protomyces spp. mating protein sequences were collected from genome assemblies and annotations with mating genes/proteins of *S. pombe* and *Pneumocystis* spp. as query. Alignment of protein sequences were acheived by ClustalX2. RAxML and rapid bootstrapping (1000x) were applied for building the trees. Bootstrap values (percent) are indicated at each node.Maps of the *matM* loci (e) and *matP* loci (f) of *Protomyces* spp. The *Protomyces* spp. contigs containing *mat* loci were applied as dnaseg in the genoplotR pakage in R. Interspecies protein sequence comparisons by tblastx with e< $e^{-100}$ and bit score ≥100 were applied as comparison in genoplotR. Coloured lines between the loci maps depicts protein comparisons in red for the same orientation and in blue for reverse orientation, where the colour depth indicates higher levels of protein similarity. Key to gene colour codes (blue: mating genes; green: conserved with *S. pombe*; yellow: conserved between most *Protomyces* spp).

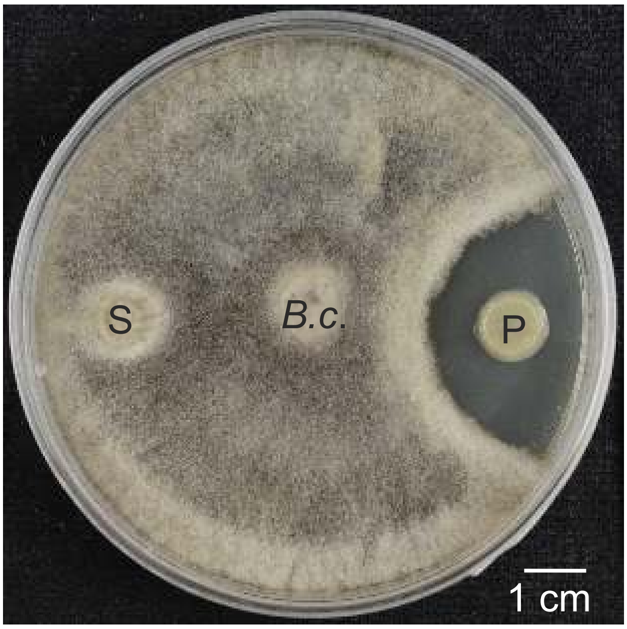


**Figure S5.** No growth inhibition effect by SC29 (S) was observed against *Botrytis cinerea* (*B.c.*). As a control, a *Paenibacillus* sp. (P) known to have antifungal activity against *B. c.* was used. GYP agar plate was used for cultivation. Photo was taken seven days post inoculation.

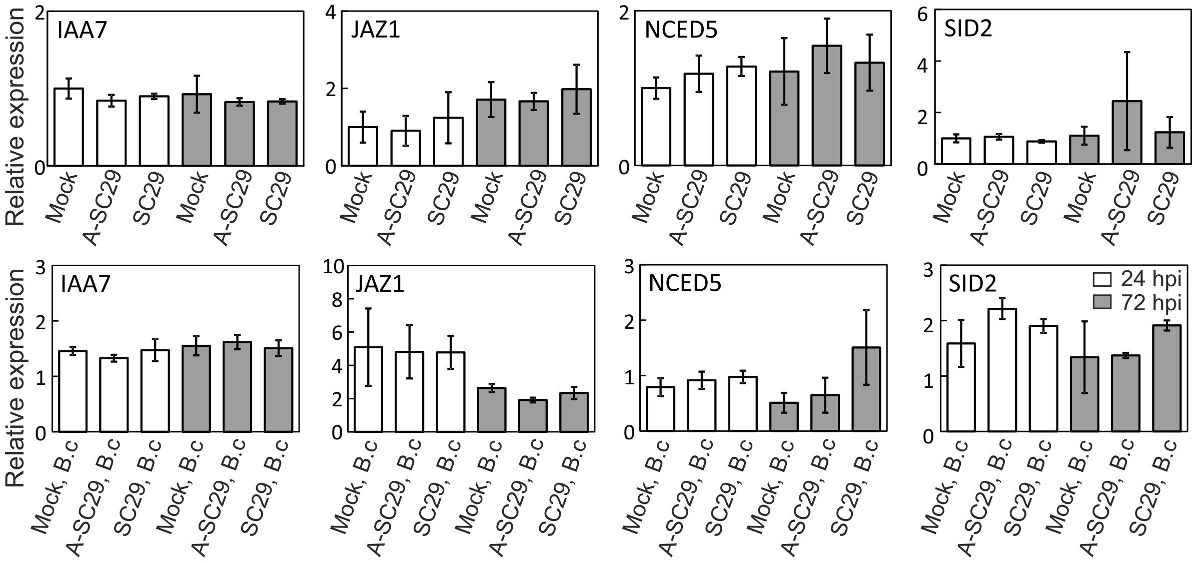


**Figure S6.** Relative gene expression of signaling genes after SC29, A-SC29, or mock treatment only (top row) or for plants pre-treated with SC29, A-SC29, or mock and subsequently infected with *Botrytis* (bottom row). Gene expression was scaled to water control 24 hrs for each gene. Data is presented as the mean ±SD of three independent biological repeats. Statistics were performed with glht package in R (3.5.1) with fdr method for adjusted p values. A-SC29: autoclave killed SC29. B.c: *Botrytis cinerea*. hpi: hour post infection. See supplementary File S4 for a list of primers, primer efficiencies, full marker gene names and their AGI codes.

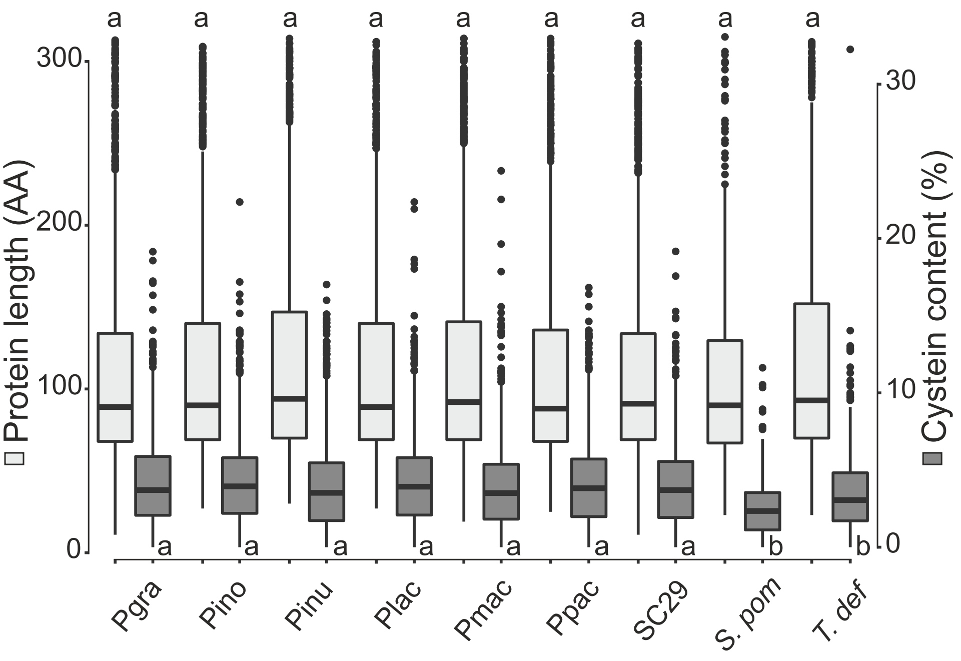


**Figure S7.** Comparison of protein length and cysteine content of short secreted proteins (SSPs) from *Protomyces* and reference species. *Protomyces* SSPs have higher average cysteine content than *Schizosaccharomyces pombe* and *Taphrina deformans*. Pgra: *P. gravidus*, Pino: *P. inouyei*, Pinu: *P. inundatus*, Plac: *P. lactucaedebilis*, Pmac: *P. macrosporus*, Ppac: *P. pachydermus*, SC29: *P.* sp. SC29. *S. pom*: *Schizosaccharomyces pombe. T. def: Taphrina deformans.* Box plots with a different letters are statistically significant from each other (P<0.05) by one-way ANOVA + Tukey HSD tests run in R.

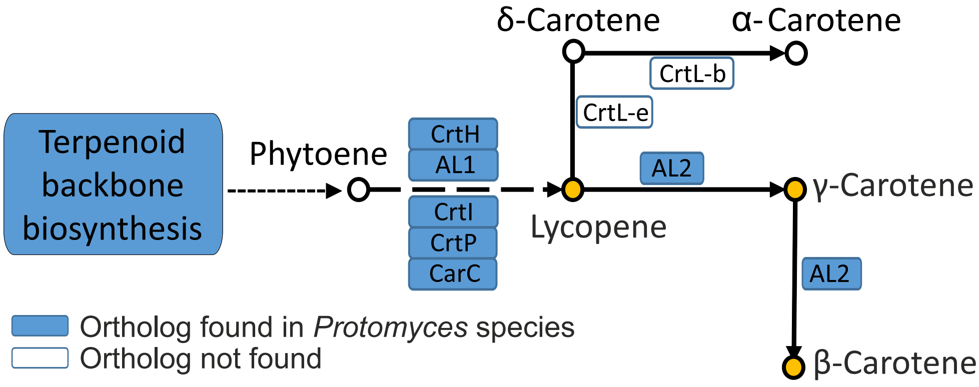


**Figure S8**. Carotenoid biosynthesis pathway in *Protomyces* species. Orthologs of known biosynthesis pathway genes that were present in all *Protomyces* species are indicated in blue boxes. Genes in white boxes were not found in any species. For sequences used in BLAST queries to identify these genes see supplemental File S10.

**Table S1.** Methods used to test *Arabidopsis* infection with SC29 yeast cells.

| Environment | Method | Concentration | Plant age | Temperature | Duration |
|---|---|---|---|---|---|
| Indoor (growth chamber) | drop | OD = 1 | 2 week | 8 °C | 21 days |
| | | OD = 1 | 2 week | 23/18 °C | 21 days |
| | | OD = 1 | 4 week | 8 °C | 7 days |
| | | OD = 1 | 4 week | 23/18 °C | 7 days |
| | spray | OD = 0.1 | 2 week | 8 °C | 21 days |
| | | | 2 week | 23/18 °C | 21 days |
| | | | 4 week | 8 °C | 7 days |
| | | | 4 week | 23/18 °C | 7 days |
| | | OD = 1 | 2 week | 8 °C | 21 days |
| | | | 2 week | 23/18 °C | 21 days |
| | | | 4 week | 8 °C | 7 days |
| | | | 4 week | 23/18 °C | 7 days |
| | infiltration | OD = 0.1 | 4 week | 8 °C | 7 days |
| | | OD = 0.1 | 4 week | 23/18 °C | 7 days |
| Outdoor | spray | OD = 0.1 | 2 week | Snow cover | ~ 7 months |
| | | OD = 1 | 2 week | Snow cover | ~ 7 months |

**Table S2**: Summary of open reading frames (Orfs) in the size range of 80-333 amino acids (AA), small secreted proteins (SSPs), and cysteine-rich SSP (CSSPs) found in the genomes of the listed Taphrinomycotina species. As a baseline reference for comparison, Orfs, SSPs, and CSSPs were identified from 13Mbp of computer generated random DNA sequence (Random sequence); the average result from five independently generated and analysed 13Mbp random sequences is shown.

| Species | Orfs (80-333 AA) | SSPs | CSSPs |
|---|---|---|---|
| *Protomyces gravidus* | 18069 | 1085 | 547 |
| *P. inouyei* | 20886 | 1376 | 718 |
| *P. inundatus* | 23865 | 1421 | 698 |
| *P. lactucaedebilis* | 20672 | 1402 | 726 |
| *P. macrosporus* | 22007 | 1235 | 615 |
| *P. pachydermus* | 19538 | 1286 | 630 |
| *P. sp.* SC29 | 18555 | 1170 | 587 |
| *Taphrina deformans* JCM 22205 | 18421 | 909 | 431 |
| *T. deformans* PYCC 5710 | 18081 | 881 | 421 |
| *T. flavorubra* JCM 22207 | 21829 | 1097 | 513 |
| *T. populina* JCM 22190 | 14424 | 799 | 378 |
| *T. wiesneri* JCM 22204 | 15314 | 756 | 334 |
| *Schizosaccharomyces pombe* | 6077 | 248 | 65 |
| *S. japonicus* | 10507 | 511 | 195 |
| *S. octosporus* | 6465 | 255 | 60 |
| Random sequence | 26526 | 198 | 53 |

**Table S3.** Enzymes involved in tryptophan (Trp)-dependent indole acetic acid (IAA) bio-synthesis pathways of *Protomyces* species. - : This enzyme is hypothetical, the gene has never been found in fungi. Species abbreviations: Pgra: *P. gravidus*, Pino: *P. inouyei*, Pinu: *P. inundatus*, Plac: *P. lactucaedebilis*, Pmac: *P. macrosporus*, Ppac: *P. pachydermus*, SC29: *P.* sp. SC29. The number of orthologous proteins found in each species is indicated for the known potential conserved tryptophan-dependent auxin biosynthesis pathways. Pathway name abbreviations are: IAM, indole-3-acetamid; IAN, indole-3-acetonitrile; IPyA, indole-3-pyruvate; TSO, tryptophan side-chain oxidase; TAM, tryptamine..* Conserved auxin efflux carrier proteins were also found in all the species. The criteria of ortholog selection are bit scores > 120, E-values > 0.05 and identities > 50%. For the sequences used in queries to find these pathways see supplementary file S11.

| Pathway | Enzymes | SC29 | Pgra | Pino | Pinu | Plac | Pmac | Ppac |
|---|---|---|---|---|---|---|---|---|
| IAM | Trp2-monooxygenase (TMO/IaaM) | 0 | 0 | 0 | 0 | 0 | 0 | 0 |
| | IAM hydrolase (IaaH) | 1 | 1 | 1 | 1 | 1 | 1 | 1 |
| IAN | unknown enzymes | - | - | - | - | - | - | - |
| | Nitrilase (NIT) | 3 | 3 | 3 | 2 | 3 | 2 | 3 |
| IPyA | Trp aminotransferase (TAM) | 1 | 1 | 1 | 1 | 1 | 1 | 1 |
| | IPyA decarboxylase (IPDC) | 1 | 1 | 1 | 1 | 1 | 1 | 1 |
| | IAAld dehydrogenase (IAD) | 2 | 2 | 2 | 2 | 2 | 2 | 2 |
| TSO | Trp side-chain oxidase (TSO) | - | - | - | - | - | - | - |
| | IAAld dehydrogenase (IAD) | 2 | 2 | 2 | 2 | 2 | 2 | 2 |
| TAM | Tryptophan decarboxylases (TDC) | 2 | 2 | 2 | 2 | 2 | 2 | 4 |
| | Amine oxidase (AOX) | 2 | 2 | 2 | 2 | 2 | 2 | 2 |
| | IAAld dehydrogenase (IAD) | 2 | 2 | 2 | 2 | 2 | 2 | 2 |
| | Flavin monooxygenase (YUC) | 0 | 0 | 0 | 0 | 0 | 0 | 0 |
| Other* | Auxin efflux carrier (1) | 1 | 1 | 1 | 1 | 1 | 1 | 1 |
| | Auxin efflux carrier (2) | 1 | 1 | 1 | 1 | 1 | 1 | 1 |

**Table S4.** Number of HHKs (hybrid histidine kinases) found in *Protomyces* species. HHK protein sequences of plant pathogenic Ascomycota fungi were applied as search queries (see file S8 for sequences used) using an E-value cutoff of $1e^{-30}$. *: Gene expansion predicted by badirate shows SC29 has five group I HHKs; one gene that only possessed a kinase domain, but lacking other characteristic HHK domains, was removed after manual curation. - : HHK was not found. Species abbreviations: Pgra: *P. gravidus*, Pino: *P. inouyei*, Pinu: *P. inundatus*, Plac: *P. lactucaedebilis*, Pmac: *P. macrosporus*, Ppac: *P. pachydermus*, SC29: *P.* sp. SC29.

| HHK (group) | SC29 | Pgra | Pino | Pinu | Plac | Pmac | Ppac |
|---|---|---|---|---|---|---|---|
| I | 4* | 1 | 1 | 1 | 1 | 1 | 1 |
| II | - | - | - | - | - | - | - |
| III | 2 | 1 | 2 | 1 | 2 | 1 | 1 |
| IV | - | - | - | - | - | - | - |
| V | 1 | 1 | 1 | 1 | 1 | 1 | 1 |
| VI | - | - | - | - | - | - | - |
| VII | - | - | - | - | - | - | - |
| VIII | 1 | 1 | 1 | 1 | 1 | 1 | 1 |
| IX | 3 | 2 | 2 | 2 | 2 | 2 | 2 |
| X | 1 | 1 | 1 | 1 | 1 | 1 | 1 |
| XI | 3 | 2 | 2 | 3 | 2 | 1 | 2 |
| Dual HK | - | - | - | - | - | 1 | - |
| Total | 15 | 9 | 10 | 10 | 10 | 9 | 10 |
